# A complex containing RhoBTB3-SHIP164-Vps26B promotes the biogenesis of early endosome buds at Golgi-endosome contacts

**DOI:** 10.1101/2022.09.30.510409

**Authors:** Jingru Wang, Qingzhu Chu, Weiping Chang, Lin Deng, Wei-Ke Ji

## Abstract

Early endosomes (EEs) are central hubs for cargo sorting in vesicular trafficking. Cargoes destined for degradation retain in EEs that are eventually delivered to lysosomes, while recycled cargo destined for the plasma membrane (PM) or to the Golgi are segregated into EE buds, a specialized membrane structure transiently generated on EEs during cargo sorting. Until now, the molecular basis of the membrane expansion during the biogenesis of EE buds is completely elusive. Here, we identify a protein complex containing a Vps13 domain-containing lipid transporter SHIP164, ATPase RhoBTB3 and retromer component Vps26B, promotes the membrane expansion in the biogenesis of EE buds at Golgi-EE contacts. SHIP164 depletion specifically reduces the size and number of EE buds marked by Vps26B and actin, and results in less but enlarged EEs, which can be substantially rescued by wild type SHIP164 other than lipid transfer-defective mutants, suggesting a role of lipid transfer of SHIP164 in the process. SHIP164 and RhoBTB3 interact with enzymes for phospholipid synthesis on motile Golgi vesicles, that frequently contact EEs. Functionally, SHIP164 depletion substantially reduces the trafficking of sphingomyelin to the PM, and impairs cell growth. Together, we propose a lipid transport-dependent route from the Golgi to EEs at the contacts required for EE bud biogenesis.

## Introduction

Although vesicular trafficking mediates the bulk transport of many lipids, mounting evidence support that lipid transporters-mediated lipid exchange at membrane contact sites (MCSs) is the major transport route for certain lipid types in specific cellular contexts^1–, 5^, including organelle biogenesis^6–8^, organelle trafficking and division^9^, and alleviation of lipotoxicity^10, 11^.

MCSs are defined as cytosolic gaps of 10 – 30 nm between one and the other organelles^1, 12^. Intracellular lipid transporters transfer lipids across opposing membranes at MCSs through two modes of action – shuttling or bridging^4, 5, 13^. Shuttle transporters typically extract one or two lipid molecules from the membrane of the donor organelle, solubilize it during transport through the cytosol and deposit it in the acceptor organelle membrane, while bridge transporters feature an extended channel, most likely lined with hydrophobic residues that bind tens of lipids at once^5, 14–16^. While a complex of synaptotagmin-like mitochondrial-lipid-binding domain-containing lipid transporters are suggested to act as shuttle transporters for glycerophospholipids and/or ceramides across MCSs in yeast and metazoen^11, 17–21^, the chorein family of lipid transporters, including the Vps13 proteins and the autophagy-related protein Atg2, that harbor an extended hydrophobic groove at their N-terminal regions (NT), bind tens of lipids at once and show much higher lipid extracting activities than shuttle transporters^15, 16, 22–24^. These bridge lipid transporters are cytosolic proteins that are recruited to MCSs via two adaptors resident on two opposing membrane of organelles. For instance, mammallian Vps13 proteins are recurited to the ER via the recognition of FFAT or phospho-FFAT motifs at their NTs by vesicle-associated membrane proteins (VAPs)^25^ or motile sperm domain containing protein 2 (MOSPD2)^26^, meanwhile these proteins are recruited to the other organelles by binding to other adaptors via their C-terminal regions^16, 22, 27^. Emerging evidence showed that the bridge lipid transporters mediate lipid transfer to meet the requirments for the quick growth of certain organelles^7^. Atg2 mediates the transport of phospholipids from the ER to growing autophagosomes upon starvation^23, 28^. Vps13D is suggested to be required for the growth of peroxisomes^29^.

Syntaxin 6-interacting protein with 164 kDa (SHIP164, also known as BLTP3B) is identified in a large (>700 kDa) complex in a pioneering study^30^. SHIP164 has a Vps13/Chorein domain at its NT, homologous with Vps13 proteins, indicating that SHIP164 is another bridge lipid transporter that is implicated in endosomes to Golgi trafficking^31^. However, the molecualr mechanism of SHIP164 localization and cellular functions remain largely elusive. In this study, we identified an ATPase RhoBTB3 on the Golgi apparatus and Vps26B, a component of retromer on early endosomes (EE), serving as adaptors to recruit SHIP164 to the Golgi-EE contacts. Functionally, lipid transfer by SHIP164 is required for the biogenesis of EE buds rather than EEs. SHIP164 associated with glycerophospholipids and sphingolipids. SHIP164 and RhoBTB3 interact with enzymes for phospholipid synthesis on motile Golgi vesicles, which are phospholipid-synthesizing/distributing organelles that may supply phospholipids for EE bud growth.

## Results

### Identification of an ATPase RhoBTB3 as a SHIP164 adaptor on the Golgi complex

Both endogenous SHIP164 and GFP tagged SHIP164 (GFP-SHIP164) are mainly cytosolic without strong membrane associations^30^ (Fig. S1A). We reasoned that the lipid transporter SHIP164 might be recruited to a specific MCS by unknown protein adaptors. To identify the adaptor, we conducted a small GTPase/ATPase library screening, in which GFP-SHIP164 was co-expressed with small GTPase/ATPase one by one, and examined the co-localization between GFP-SHIP164 and each small GTPase/ATPase in HEK293 cells by live-cell confocal microscopy (Fig. S1A). We used HEK293 cells for this purpose, as these cells were more suitable for the transfection and expression of large SHIP164 constructs (∼10k bp). To this end, we identified RhoBTB3, an ATPase resident on the Golgi complex^32^, as an adaptor for SHIP164 (Figs. 1A & S1B), as revealed by a strong recruitment of GFP-SHIP164 to Halo-RhoBTB3-postive membranes in both the peri- nuclear and cell periphery by line-scan analyses. We then used 3-dimension (3D) rendering of z-stacks to examine the localizations of GFP-SHIP164 and Halo-RhoBTB3 relative to the Golgi apparatus. Both GFP-SHIP164 and Halo-RhoBTB3 were specifically enriched on the Golgi complex in 3D (Fig. 1B), as revealed by co-localization analysis based on x-z and y-z projections of 3D rendering. Consistent with the imaging results, an interaction between GFP-SHIP164 and Halo-RhoBTB3 was confirmed by GFP-Trap assays (Fig. 1C). Consistently, we found that Halo-RhoBTB3 is mainly on the cis/medial Golgi ribbon with a significant portion of Halo-RhoBTB3 associated with trans-Golgi vesicles (Fig. S2A).

**Fig. 1.**
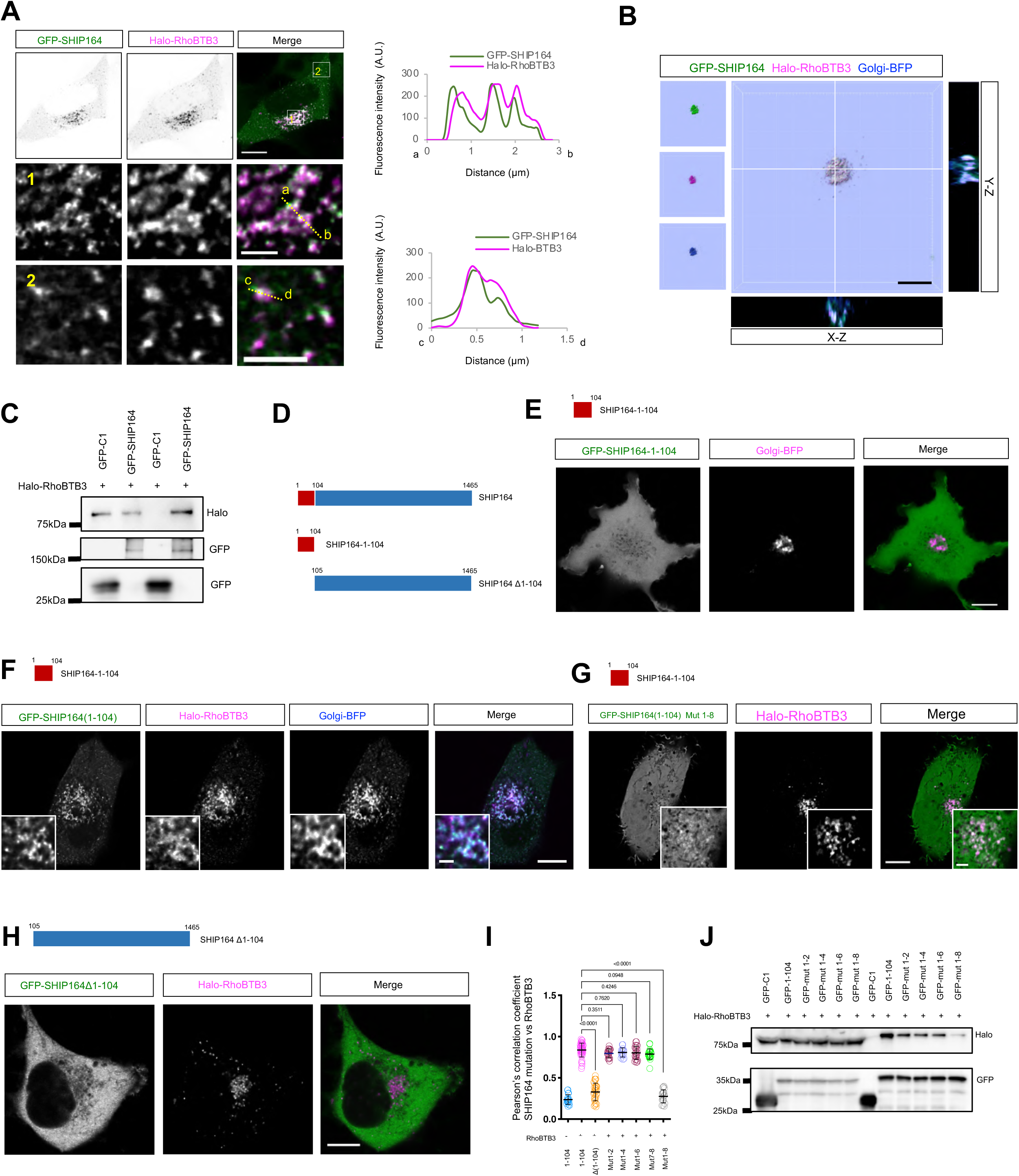
Identification of ATPase RhoBTB3 as a SHIP164 adaptor on the Golgi. **A.** Representative images of a live HEK293 cell expressing GFP-SHIP164 (green) and Halo-RhoBTB3 (magenta). Two boxed regions were shown at the bottom with line-scan analyses on the right. **B.** Representative 3D rendering of a HEK293 cell expressing GFP-SHIP164 (green), Halo- RhoBTB3 (magenta), and Golgi-BFP (B4GALT1) (blue) with y-z projection to the right and x-z projection to the bottom. **C.** GFP-Trap assays demonstrate an interaction between GFP-SHIP164 and Halo- RhoBTB3 in HEK293 cells. **D.** Diagrams of two SHIP164 truncation mutants. **E, F**. Representative images of a HEK293 cells expressing GFP-SHIP164-1-104 (green) and Golgi-BFP without (**E**) or with Halo-RhoBTB3 (**F**) with insets. **G.** Representative images of a HEK293 cells expressing GFP-SHIP164-mut1-8(green) and Halo-RhoBTB3 (magenta) with insets. **H.** Representative images of a HEK293 cells expressing GFP-SHIP164- Δ1-104 (green) and Halo-RhoBTB3 (magenta). **I.** Pearson’s correlation coefficient of SHIP164 proteins vs the Golgi; GFP-SHIP164-1-104 without Halo-RhoBTB3 (67 cells); GFP-SHIP164- Δ1-104 (29 cells); GFP-SHIP164-1-104- Mut1-2 (18 cells); GFP-SHIP164-1-104-Mut1-4 (15 cells); GFP-SHIP164-1-104-Mut1-6 (23 cells); GFP-SHIP164-1-104-Mut7-8 (22 cells); or GFP-SHIP164-1-104-Mut1-8 (19 cells) in more than 3 independent experiments. Ordinary one-way ANOVA with Tukey’s multiple comparisons test. Mean ± SD. **J.** GFP-Trap assays demonstrate interactions between GFP-SHIP164 mutants and Halo- RhoBTB3 in HEK293 cells. Scale bar, 10μm in the whole cell images and 2μm in the insets in (A, B, E, F, G & H).

### A N-terminal region of SHIP164 is responsible for interacting with RhoBTB3

We sought to explore the mechanism underlying SHIP164 recruitment by RhoBTB3. Through the dissection of SHIP164 (Fig. 1D), we found that a N-terminal region containing residues 1–104 (SHIP164-1-104) was cytosolic in absence of Halo-RhoBTB3 (Fig. 1E), but was strongly recruited to RhoBTB3-postive Golgi complex (Fig. 1F). Further, SHIP164 with a deletion of this region (SHIP164-Δ1-104) failed to be recruited by Halo-RhoBTB3 (Fig. 1G), indicating that the residues 1-104 of SHIP164 is responsible for its recruitment to the Golgi complex by RhoBTB3. We referred this region as to RhoBTB3-interacting domain (RBID) thereafter in this manuscript.

Further, we found that 8 hydrophobic residues in the RBID, which were highly conserved among the Vps13 domain-containing proteins (Fig. S2B; highlighted in yellow) were required for its interaction with RhoBTB3, and substitutions of these hydrophobic residues to hydrophilic or non-polar resides substantially reduces the recruitment, as revealed by live cell microscopy (Figs. S2C-F; & 1G) and GFP-Trap assays (Fig.1J).

RhoBTB3 was reported to be an ATPase of Rho GTPase family that is required at endosome to Golgi transport, and it contained a putative Rho domain and a BTB domain with multiple point mutations^32^ (Fig. S3A). To explore how RhoBTB3 interacted with SHIP164, we examined the recruitment of the RBID of SHIP164 by RhoBTB3 mutants by live-cell confocal microscopy and GFP-Trap assays. RhoBTB3-N138D, in which the binding and hydrolysis of ATP was impaired^32^, still recruited SHIP164-RBID. Among 3 mutants, A498T, D532E, and I533K, in which the interactions of RhoBTB3 with Rab9 were hampered^32^, The D532E mutation significantly reduced the RhoBTB3-mediated recruitment of the SHIP164-RBID, whereas the other 2 mutants were still able to mediate the recruitment (Fig. S3B-E).

In addition, we found that SHIP164 may self-associate via the RBID (Fig. S4A, B), and GFP-Trap assays indicated that RhoBTB3 may also self-associate via multiple regions (Fig. S4C-G). These results may explain, in part, the fact that SHIP164 is originally identified in the large (>700 kDa) complex^30^.

### Identification of Vps26B as another adaptor of SHIP164 on EEs

Our results suggest that SHIP164 likely functions at Golgi-associated MCSs. Given that this bridge lipid transporter used its RBID at the NT to target the Golgi complex, we reason that SHIP164 may recognize another adaptor on the other organelles via its CT. Then, we sought to identify another adaptor by GST-pulldown assays followed by mass spectrometry (MS) using purified GST-SHIP164-CT (C-terminal region containing residues 890-1465) as a bait in mouse brain lysates, in which SHIP164 was highly expressed (Fig. 2A-C). After removal of proteins co-pelleted by GST tag alone, we found that a couple of proteins that are functionally associated with the Golgi-endosome trafficking (Fig. S5A). We then explored the relationships between SHIP164-CT and these protein candidates by live-cell confocal microscopy (Fig. S5B-E). To this end, we identified Vps26B, a component of retromer, functioning at the retrograde transport from endosomes to the Golgi^33^, as another adaptor for SHIP164. Importantly, Vps26B, but not its paralog Vps26A, substantially recruited SHIP164-CT (Fig. 2D, E). Consistently, GFP-Trap assays showed that Halo- Vps26B, but not Halo-Vps26A, interacted with GFP-SHIP164-CT (Fig. 2F), suggesting that Vps26B and Vps26A are paralogs, in agreement with a previous study^34^. In addition, Vps26B could recruited the full-length SHIP164 (Fig. 2G).

**Fig. 2.**
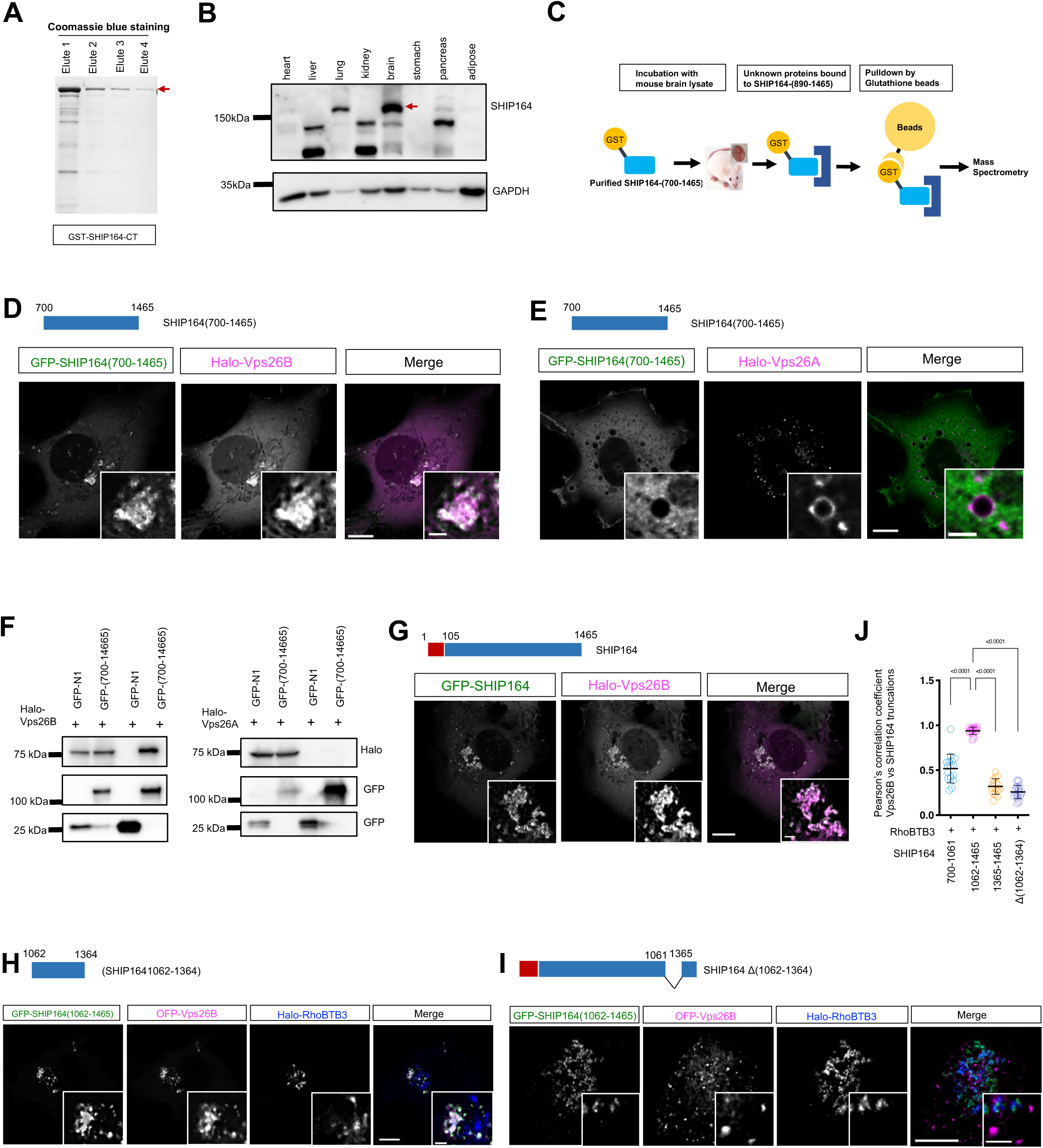
Vps26B recruits SHIP164 to early endosomes. **A.** Coomassie blue staining of purified GST-SHIP164-CT. **B.** Western blots of endogenous SHIP164 in mouse tissues. Red arrow denoted SHIP164. **C.** Schematic cartoon of GST pulldown assays using purified SHIP164-CT in mouse brain. **D, E**. Representative images of HEK293 cells expressing GFP-SHIP164-CT (green), along with either Halo-Vps26B (magenta; **D**) or Halo-Vps26A (magenta; **E**) with insets. **F.** GFP-Trap assays demonstrate interactions between GFP-SHIP164-CT and Halo- Vps26B, but not Halo-Vps26A, in HEK293 cells. **G.** Representative images of a HEK293 cell expressing GFP-SHIP164 (green) and Halo- Vps26B (magenta) with insets. **H, I**. Representative images of a HEK293 cell expressing either GFP-SHIP164 (1062-1364) (green; **H**) or SHIP164 Δ (1062-1364) (green; **I**) along with OFP-Vps26B (magenta) and Halo-RhoBTB3 (blue) with insets. **J**. Pearson’s correlation coefficient of Vps26B vs either GFP-SHIP164-700-1061 (17 cells), GFP-SHIP164-1062-1465 (21 cells), GFP-SHIP164-1365-1465 (16 cells), or GFP- SHIP164- Δ (1062-1465) (23 cells) in more than 3 independent experiments. Ordinary one-way ANOVA with Tukey’s multiple comparisons test. Mean ± SD. Scale bar, 10μm in the whole cell images and 2μm in the insets in (D, E, G, H&I).

Next, we sought to understand the mechanism underlying the recruitments of SHIP164 by Vps26B through dissections of Vps26B and SHIP164-CT (Fig. S6A). Live-cell microscopy showed that the NT (residues 1-160; Fig. S6B, D), but not the CT of Vps26B (Fig. S6C, D), was able to recruit SHIP164-CT to EEs. Meanwhile, a region of SHIP164-CT (SHIP164- 1062-1364) was recruited by Vps26B, but was not recruited by RhoBTB3 (Fig. 2H, J). In contrast, the remaining portions of ship164-CT (SHIP164-700-1061 or 1365-1645) failed to be recruited by OFP-Vps26B (Fig. S6E, F). Consistently, SHIP164 with a deletion of the region containing the residues 1062-1364 (SHIP164-Δ-1062-1364) failed to be recruited by Vps26B-OFP, but was recruited by Halo-RhoBTB3 (Fig. 2I, J). Together, these results indicate that Vps26B acts as another adaptor to recruit SHIP164 to EEs by recognizing a CT region of SHIP164.

### SHIP164, Vps26B, and RhoBTB3 forms a tethering complex at Golgi-endosome contacts

The recruitment of SHIP164 to the Golgi and EE via two distinct and distal regions suggested a tetering role of SHIP164 in mediating a novel type of MCSs between the Golgi and EEs. To begin with, we explored the spatial relationship between the Golgi and EEs in immunofluoresnce staining (IF) in human retina pigment epithelial cells (RPE1). We used RPE1 cell for functional studies as this cell line featured a near-diploid karyotype with a modal chromosome number of 46 and lacks transformed phenotypes. In control cells, the Golgi (anti-GM130) and EEs (anti-EEA1) closely associated with each other, as revealed by a large number of EEs adjacent to the Golgi. The specificity of EEA1 antibody in IF was verified by siRNA-mediated depletion (Fig. S7A, B). SHIP164 depletion by two distinct siRNAs consistently showed that Golgi-EE association were not substantially reduced, and a significant number of EE still associated with the Golgi (Fig. 3A, B). However, the Golgi- EE association observed in IF from confocal microscopy may not indicate the existence of bona fide MCSs. Therefore, we conducted in situ proximity ligation assays to quantitatively examine the role of SHIP164 at this contact. Contrary to PLA-negative controls, in which only one primary antibody was used (Fig. 3C, D), the Golgi and EEs indeed formed extensive contacts in scrambled siRNA treated cells in presence of anti-GM130 and anti- EEA1, as revealed by a large number of PLA puncta. Importantly, SHIP164 depletion resulted in a ∼3-fold decrease in the number of PLA puncta, suggesting an important role of SHIP164 in the formation or maintenance of Golgi-EE contacts. Interestingly, we found that SHIP164 depletion caused less but enlarged EEs (Fig. 3A), which may also contribute to such a dramatic reduction in Golgi-EE contacts in SHIP164-depleted cells.

**Fig. 3.**
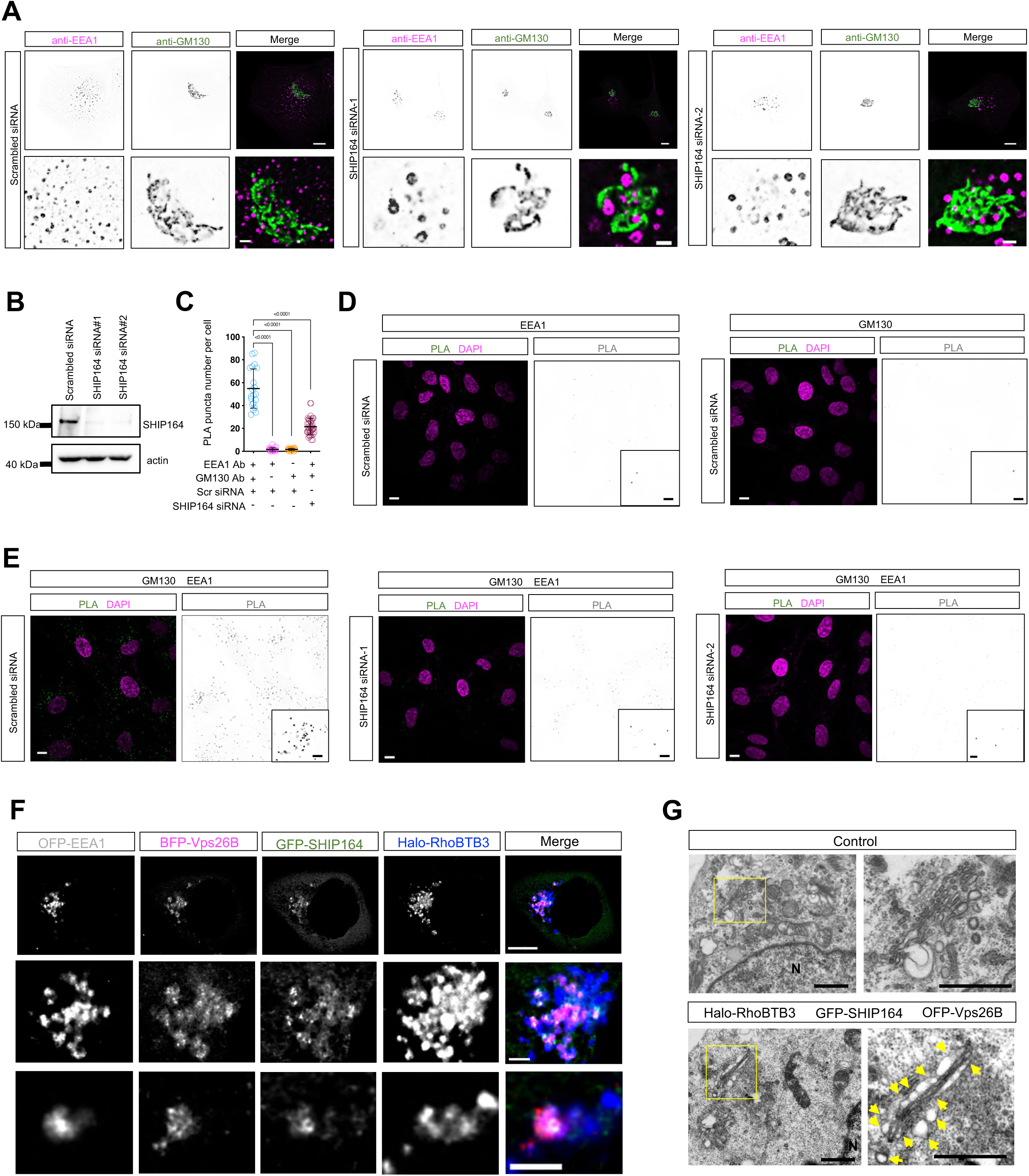
SHIP164 is required for the physical interactions between the Golgi and EE. **A.** Representative images of fixed RPE1 cells labeling endogenous EEA1 (magenta) and GM130 (green) with insets on the bottom were treated with either scrambled (left) or two SHIP164 siRNAs (middle and right). **B.** Immunoblots showing the efficiency of siRNA-mediated depletion of SHIP164. **C-E**. In situ proximity ligation assays to monitor the effects of SHIP164 on Golgi-early endosome associations in fixed RPE1 cells probed with primary antibodies (anti-EEA1 for EE; anti-GM130 for the Golgi) followed by secondary antibodies coupled to specific oligonucleotides. EEA1 only (20 cells) or GM130 only (20 cells) were used as negative controls; 20 scrambled cells and 40 SHIP164 siRNA-treated cells were quantified from 3 independent experiments. Ordinary one-way ANOVA with Tukey’s multiple comparisons test. Mean ± SD. **F.** Representative images of a HEK293 cell expressing GFP-SHIP164 (green), Halo-RhoBTB3 (blue), BFP-Vps26B (magenta) and OFP-EEA1 (gray) with insets on the bottom. **G.** Representative TEM micrographs of control (top panel) and HEK293 cell expressing GFP-SHIP164, OFP-Vps26B, and Halo-RhoBTB3 with boxed regions on the right. Yellow arrows denoted vesicular structures closely associating with the Golgi. Scale bar, 10μm in the whole cell images and 2μm in the insets in (A, D-F) and 10μm in (G).

The golden stardard for a tether is to able to increase the affinity of one to the other organelle at contacts. Therefore, we further verified the tethering ability of SHIP164 at Golgi-EE contacts by over-expression. Remarkably, overexpression of SHIP164 with its two adaptors RhoBTB3 and Vps26B resulted in a strikingly tight association between the Golgi and EEs, and almost all EEs were adjacent to the Golgi with GFP-SHIP164 specifically enriching at these contact sites (Fig. 3F).

Further, we directly examined the interactions between the Golgi and EEs by transmission electron microscopy (TEM). Electron micrographs revealed that uncoated vesicular structures, likely endosomes, extensively contacted both faces of Golgi stacks, upon RhoBTB3-SHIP164-Vps26B overexpression in HEK293 cells (Fig. 3G), which was rarely observed in control cells.

In addition, we examined the tethering ability of RhoBTB3-SHIP164-Vps26B by knocksideways assays. Taking advantage of interactions between RFP and RFP nanobody (RFPnb), we tagged Vps26B with RFPnb, and targeted Vps26B-RFPnb to the membranes of various organelles including the ER (mCh-Sec61β), the OMM (TOM20- mCh), lysosome (Lamp1-mCh), and lipid droplet membranes (ACSL3-mCh). We confirmed that the VPS26B-RFPnb was efficiently recruited to the ER via mCh-Sec61β (Fig. S8A). Remarkably, these membranes were strongly recruited to Halo-RhoBTB3-positive Golgi with specific enrichments of GFP-SHIP164 at the interfaces (Fig.S8B-F). Together, these results suggested that RhoBTB3-SHIP164-Vps26B may act as a structural tether at Golgi- EE contacts.

### Lipid transfer of SHIP164 promotes EE bud growth

Bridge lipid transporters, including Vps13 proteins and Atg2, were responsible for the biogenesis of certain organelles^23, 28, 29, 35–37^. The bridge transporter SHIP164 localizes to Golgi-EE contacts, prompting us to explore whether SHIP164 is required for the biogenesis of these two organelles. We found that the number or size of the trans-Golgi was not substantially affected upon SHIP164 depletion (Fig. S9A), though the morphology of cis-Golgi appeared to be slightly changed (Fig. 3A), suggesting that SHIP164 may not play a major role in biogensis of the Golgi. In contrast, SHIP164 depletion caused a robust reduction in number but increase in size of EEA1-labeled EEs (Fig. 3A), suggesting a potential role of SHIP164 on EE homeostasis.

Further, we sought to confirm the role of SHIP164 on EE homeostasis. Due to the heterogeneity of endosomes, we examined the effects of SHIP164 depletion on different endosomal subpopulations marked by distinct Rab proteins. We found that SHIP164 depletion had strong effects on Rab4 or Rab14-marked EEs, with Rab14-marked EE being the most affected, compared with other Rabs, including Rab5 and Rab7 (Fig. S9B-J). Therefore, we focused on the Rab14-labeled EE hereafter.

EEs are central hubs for cargo sorting, and recycled cargo destined for the plasma membrane (PM) or to the Golgi are segregated into endosome buds, a specialized membrane structure transiently generated on EEs during cargo sorting, while cargoes destined for degradation at lysosomes remain in EEs. Microtubules and their motor proteins, branched actin networks generated by the Arp2/3 activator WASH, the retromer, and structural membrane shaping proteins such as sorting nexins, have all been implicated in endosome bud structures and cargo sorting^38–41^(Fig. 4A).

**Fig. 4.**
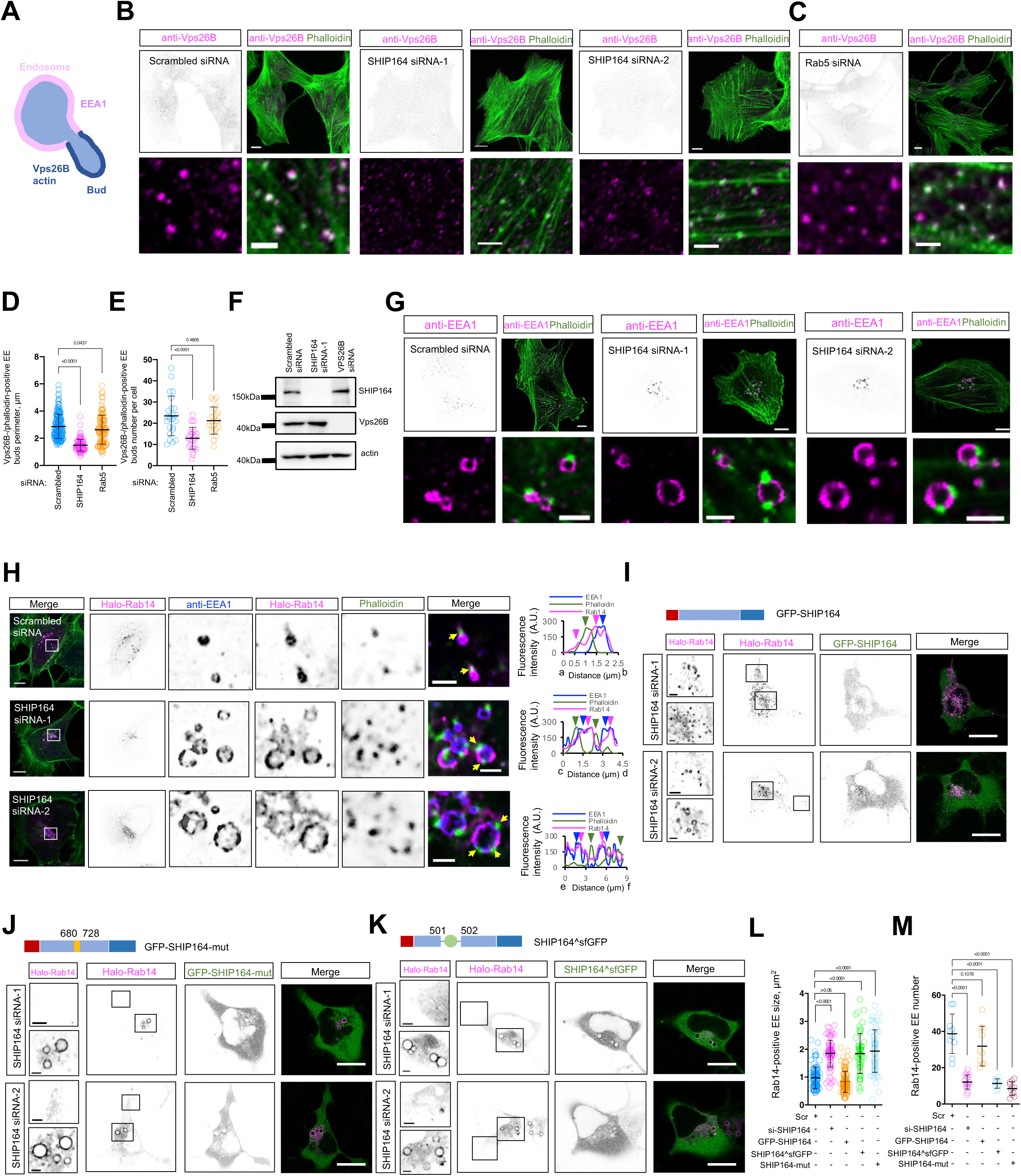
SHIP164 is required for the growth of early endosome buds. **A**. Schematic cartoon showing two EE structures, EE and an EE bud. **B, C**. Representative images of fixed RPE1 cells probed with antibodies against endogenous Vps26B (magenta) and phalloidin (green) upon scrambled (left; **B**), SHIP164 siRNA (right; **B**), or Rab5 siRNA (**C**) with insets on the bottom. **D, E**. The size (**D**) and number (**E**) of endosome buds positive for both Vps26B and actin in cells. More than 20 cells were quantified for each condition from 3 independent experiments. Ordinary one-way ANOVA with Tukey’s multiple comparisons test. Mean ± SD. **F.** Immunoblots showing that SHIP164 depletion did not affect the level of Vps26B, or vice versa. **G.** Representative images of fixed RPE1 cells probed with antibodies against endogenous EEA1 (magenta) and phalloidin (green) upon scrambled or SHIP164 siRNA with insets on the bottom. **H.** Representative images of fixed RPE1 cells expressing Halo-Rab14 (magenta) and stained with antibodies against endogenous EEA1 (blue) and phalloidin (green) with insets and line-scan analyses on the right. **I-K**. Representative images of SHIP164-depeleted HEK293 cells expressing siRNA-resistant GFP-SHIP164 (**I**), GFP-SHIP164-mut (**J**), or SHIP164^sfGFP (**K**), along with Halo-Rab14 (magenta) with insets. **L, M**. The perimeter (**L**) and number (**M**) of endosomes in (**I**-**K**). More than 20 cells were quantified for each condition from 3 independent experiments. Ordinary one-way ANOVA with Tukey’s multiple comparisons test. Mean ± SD. Scale bar, 10μm in the whole cell images and 2μm in the insets in (B, C, G, & H-K).

Vps26B is an essential retromer component distinct from Vps26A that functions at endosome buds^34^. Indeed, the IF assays showed that endogenous Vps26B formed foci, and many of these foci coloclaized with actin filaments (labeled by phalloidin), presumbly representing active sites for cargo sorting, i.e., endosome buds (Fig.4B). The specificity of Vps26B antibody used in IF was verified by siRNA-mediated depletion (Fig. S10). Intriguingly, we found that depletion of SHIP164, but not Rab5, substantially reduced the size and number of EE buds marked by endogenous Vps26B and actin (Fig. 4B-E), suggesting a potential role of SHIP164 in the formation of EE buds. Importantly, the effect of SHIP164 depletion on EE buds were not due to the decreased level of Vps26B, as immunoblots showing a similar level of Vps26B between control and SHIP164-depelted cells (Fig. 4F).

Given that actin polymerization specifically occurs at the base of endosome buds^41^, a key step of cargo sorting, we then examined the spatial relationship between EEs (anti-EEA1) and actin filaments (Phalloidin). Notably, contrary to Vps26B, EEA1 exclusively decorates EEs, but not EE buds, as revealed by a substantially lower colocalization between EEA1 and actin (Fig. 4G). In control cells, each sorting-active EE typically has one actin foci, representing one bud base on one EE (Fig. 4G). Strikingly, SHIP164 depletion resulted in enlarged EEs with two or more actin foci (Fig. 4G). Consistently, an enlarged EE also had two or more Coronin1C foci, a key regulator of actin polymerization at the base of endosome buds^41^, in SHIP164-depleted cells (Fig. S11).

Notably, contrary to EEA1 that exclusively labeled EEs, Rab14 localizes to both EEs and EE buds, with actin foci spefcifically resided at the bud base (Fig. 4H). Importantly, upon SHIP164 depletion, the buds marked by Rab14 were completely disapppeared on these enlarged EEs, while actin foci were still present on EEs (Fig. 4H), further supporting the essential role of SHIP164 in EE bud formation, and suggested that SHIP164 functions downstream of actin polymerization in EE bud formation.

Although our results showed that SHIP164 depletion strongly affected the homeostasis of EE, with a reduction in number but increase in size, it is unclear whether SHIP164 depletion affected the biogenesis of EE. We found that depletion of Rab5 significantly rescued the EE phenotypes in SHIP164-depleted cells, with the number or size of EEA1-labeled EE being restored to be normal (Fig. S12A-G). Meanwhile, depletion of either Rab5 or EEA1, or expression of dominant negative Rab5-S34N also restored the abnormal size/number of Rab14-labeled EE resulted from SHIP164 depletion (Fig. S12H-N). These results suggest that SHIP164 depletion may affect EE fusion as a consequence of the impaired EE bud biogenesis, but may not directly affect the EE biogenesis.

The defect in the formation of Rab14-labeled EE buds was specific to SHIP164, since introduction of WT-SHIP164 could almost completely rescue the phenotype (Fig. 4I). Next, we investigate whether and to what extent the EE bud defect was depedent on the lipid transfer activity of SHIP164. For this purpose, we made two lipid transfer-deficient mutants. In the first mutant (Mut-1), a couple of conserved hydrophobic residues in the midway of the hydrophobic groove of SHIP164 were mutated to hydrophilic residues to block lipid transport according to a recent study on Vps13 (Fig. S13A, B)^15^. In the second mutant (Mut-2), a superfolder GFP was inserted in the midway of hydrophobic groove (at residue 501) with GSSGSS linkers at each side of sfGFP, which hampered the movements of lipids through the groove. Importantly, the two lipid transfer-deficient mutants failed to rescue the defect in EE bud biogenesis resulted from SHIP164 depletion (Fig. 4J, K). The striking difference between WT and lipid transfer-deficient mutants in rescue experiments were not due to their expression levels, as immunoblots showing a similar level of these two proteins (Fig. S13C). These results suggest that the lipid transfer activity of SHIP164 is required for EE bud biogenesis.

Next, we sought to identify lipid species bound by GFP-SHIP164 in HEK293 cells. Lipid species associating with GFP-SHIP164 were examined by non-targeted lipidomics using liquid chromatography-tandem mass spectrometry (LC-MS/MS) with rigorous washes of the protein before lipid analysis, according to the protocol used in our previous study (Fig. 5A)^21^. GFP tag-only was set up as a negative control in the assay. GFP-SHIP164 substantially associated with glycerophospholipids, inlcuding phosphotidylcholine (PC) (∼35%), phosphatidylethanolamine (PE) (∼20%), phosphatidylserine (PS) (∼18%), and phosphoinositides (PI) (∼5%) (Fig. 5B), in agreements with recent studies on Vps13 proteins^22, 31^. In contrast, GFP-SHIP164-RBID failed to substantially associate with lipids (Fig. 5C, D), suggesting that the hydrophobic residues of the RBID may be involved in the interaction with RhoBTB3 but not lipid binding. In addition, we found that GFP-SHIP164 could also bind sphingolipids [SM and Ceramide (Cer)] (∼10%) (Fig. 5D). Taking into consideration the lipid composition of total membranes in HEK293 cells, glycerophospholipids accounted for >70% of the total phsopholipids, whereas sphingolipids only accounts for less than 15% (Fig. 5B)^21^, it is plausible that GFP-SHIP164 may also efficiently associate with sphingolipids in addition to glycerophospholipids. It should be noted that the associations between SHIP164 and sphingolipids may be indirect.

**Fig. 5.**
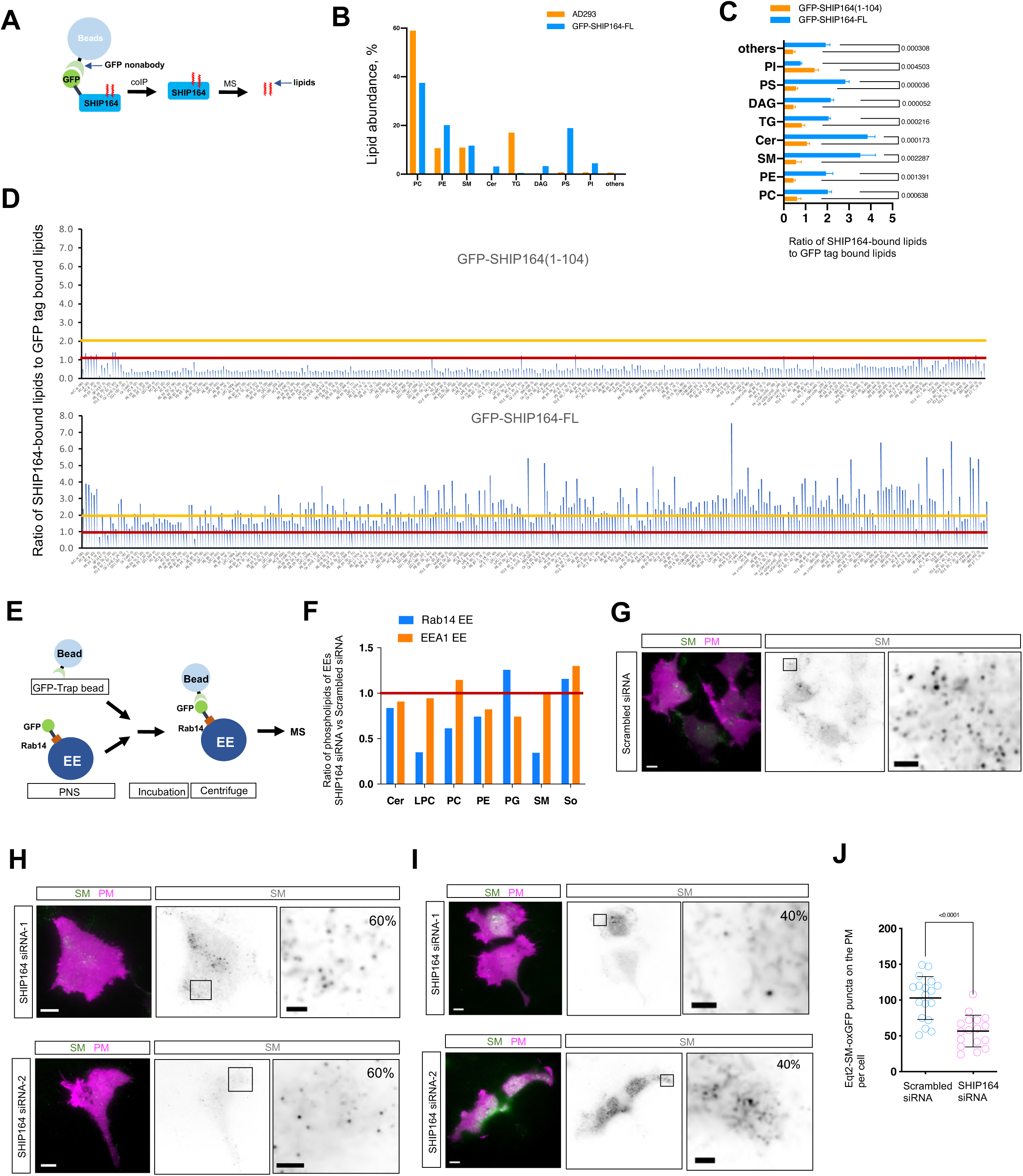
GFP-SHIP164 associates with glycerophospholipids and sphingolipids in HEK293 cells. **A.** Schematic diagram of non-targeting lipidomic analysis of GFP-SHIP164 and GFP- SHIP164-1-104. **B.** The abundance of lipids bound to GFP-SHIP164, from three independent assays. The lipid composition of total membranes in HEK293 cells were shown^21^. **C.** Ratio of phospholipid species bound by either GFP-SHIP164-1-104 or GFP-SHIP164 to GFP tag alone, from three independent assays. Two-tailed unpaired student t-test. Mean± SD. **D.** Ratio of each phospholipid specie bound by either GFP-SHIP164-1-104 (top), or GFP- SHIP164 (bottom) to GFP tag alone, from three independent assays. Red line denoted ratio=1, and yellow line indicated ratio =2. **E.** Schematic diagram of enrichments of GFP-Rab14 or GFP-EEA1 labeled EEs followed by non-targeting lipidomics. PNS: post-nuclear supernatant. **F.** The level of phospholipids of GFP-Rab14 or GFP-EEA1 labeled EEs in SHIP164 depleted HEK293 cells normalized to those in scrambled siRNA-treated cells, from three independent assays. **G-I**. Representative TIRF images of HEK293 cells expressing Eqt2-SM-oxGFP (SM sensor; green), and PLC-delta-PH-iRFP (PM marker; magenta) with insets upon scrambled (**G**) or SHIP164 siRNA (**H**, **I**). SM sensor formed puncta on the PM in ∼60% of SHIP164 depleted cells (**H**); whereas displayed ER-like structures in the 40% of SHIP164 depleted cells (**I**). **J.** The number of SM puncta per cell upon scrambled (18 cells) or SHIP164 siRNA (17 cells) in (**G, H**). Two-tailed unpaired student t-test. Mean ± SD. Scale bar, 10μm in the whole cell images and 2μm in the insets in (G-I).

Next, we examined the effects of SHIP164 depletion on lipid composition of EE membranes using non-targeted lipidomics using LC-MS/MS. We used GFP-trap assays to enrich GFP-Rab14 or GFP-EEA1-positive EEs (Fig. 5E) and examined the lipid compositions of EE membranes. We found that, in case of EEA1-labeled endosomes, SHIP164 depletion did not subtantially affected the ratio of phospholipids compared to control (Fig. 5F), though levels of some specific phospholid species are indeed altered (Fig. S13D). In contrast, SHIP164 depletion resulted in a significant reduction in levels of LPC/PC and SM, but did not strongly affect the levels of other phospholipids (Cer, PE, PG, or So) on the membranes of Rab14-labeled EEs (Figs. 5F & 13E), suggesting that SHIP164 may directly transfer these lipids to promote the biogenesis of EE buds.

It is intriguing to us that SHIP164 associated with and was implicated in the regulation of SM levels of EE membranes. Since SM is mainly synthesized on the Golgi, and transported to the PM via yet unclear routes, we thus explored whether SHIP164 contributed to the trafficking of SM to the PM. The SM trafficking to the PM was robustly reduced upon SHIP164 depletion (Fig. 5G-I), as revealed by a SM sensor on TIRF microscopy. Notably, in control cell, SM formed foci on the PM (Fig. 5G), and the number of SM foci was reduced by ∼2-fold in ∼60% of SHIP164 depleted cell (Fig. 5H), meanwhile in remaining ∼40% of SHIP164 depleted cells, SM signals displayed an ER-like pattern (Fig. 5I), which was rarely observed in control cells, suggesting that much of SM might be mistakenly transported to the ER instead of the PM (Fig. 5I). The question of how SHIP164 regulates SM trafficking to the PM remains unclear, and one possible explanation is that SHIP164 may directly or indirectly promote the transfer of SM from the Golgi to EEs, and part of the latter recycles back to the PM.

The sphingolipids on the PM are essential components of lipid rafts, which mediates vesical trafficking, cell polarity, and receptor-mediated cellular signaling^42^. Importantly, we observed that SHIP164 depletion robustly impaired cell growth, as revealed by a significant delay in the growth curve upon SHIP164 depletion (Fig. S14). Collectively, these lines of evidence support that SHIP164 may play a role in the trafficking of SM to the PM, and promotes cell growth, in addition to a defect in the retrograde trafficking of certain cargoes^31^.

### SHIP164 and RhoBTB3 interact with enzymes in phospholipid synthesis pathways on the Golgi

We then asked how the SHIP164-mediated lipid transfer was fullfilled in cellular contexts. The ER is the main site for the synthesis of majority of lipids^43^. Therefore, we asked whether SHIP164 could also associate with the ER by live-cell confocal microscopy. GFP- SHIP164 did not localize to the ER (Fig. S15A). Over-expression of ER adaptors, including Halo-VAPA (Fig. S15B), Halo-VAPB (Fig. S15C), or Halo-MOSPD2 (Fig. S15D), failed to recruit GFP-SHIP164 to the ER, suggesting that SHIP164 was not a typical lipid transporter associating with the ER.

The de novo synthesis of sphingolipids involves a series of enzymes, inlcuding ER-resident rate-limiting serine palmitoyltransferases (SPTLCs) and Cer synthases (CerS), and Golgi-resident SM synthases (SMS)^44^. Importantly, we found that a small but significant portion of GFP-SHIP164 associated with SPTLC3 puncta (Fig. 6A) but not with ER-resident Halo- GPAT4 (Fig. S16A) or Halo-HMGCR1 (Fig. S16B). Consistently, GFP-Trap assays showed that GFP-SHIP164 interacted with Halo-SPTLC3 in HEK293 cells (Fig. 6B). Further, RhoBTB3 strongly associated with SPTLC3 on both the Golgi (Fig. 6C), to a much higher extent than the SHIP164-SPTLC3 association (Fig. 6F). GFP-trap assays indicated that GFP-RhoBTB3 interacted with Halo-SPTLC3 as well (Fig. 6D). There results suggested that SHIP164, RhoBTB3, and SPTLC3 may form a protein complex. Indeed, GFP-SHIP164, Halo-RhoBTB3, and SPTLC3-SNAP were well co-localized, presumbly on the Golgi complex (Fig. 6E, F).

**Fig. 6.**
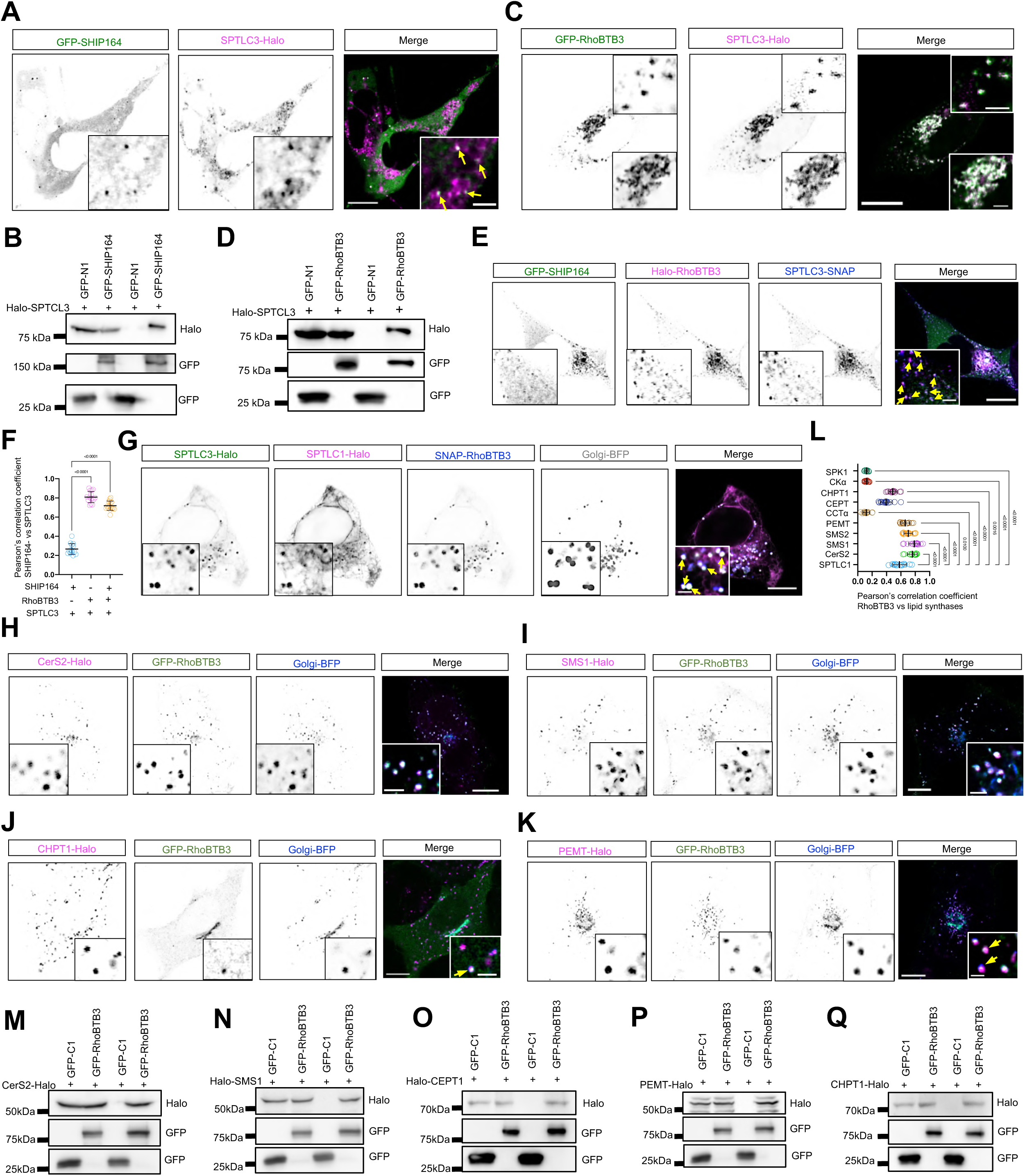
SHIP164 and RhoBTB3 interact with enzymes in phospholipid synthesis pathway on the Golgi. **A**. Representative images of a HEK293 cell expressing GFP-SHIP164 (green) and SPTLC3-Halo (magenta) with insets. Yellow arrows denoted GFP-SHIP164 associating with SPTLC3 foci. **B**. GFP-Trap assays demonstrate an interaction between GFP-SHIP164 and SPTLC3-Halo in HEK293 cells. **C**. Representative images of a HEK293 cell expressing GFP-RhoBTB3 (green) and SPTLC3-Halo (magenta) with insets. Yellow arrows denoted GFP-RhoBTB3 associating with SPTLC3 foci. **D**. GFP-Trap assays demonstrate an interaction between GFP-RhoBTB3 and SPTLC3-Halo in HEK293 cells. **E**. Representative images of a HEK293 cell expressing GFP-SHIP164 (green), Halo-RhoBTB3 (magenta) and SPTLC3-SNAP (blue) with insets. Yellow arrows denoted GFP- SHIP164 and Halo- RhoBTB3 associating with SPTLC3 foci. **F**. Pearson’s correlation coefficient of GFP- SHIP164 vs SPTLC3 in presence (14 cells) or absence of Halo-RhoBTB3 (15 cells), or RhoBTB3 vs SPTLC3 (16 cells) from 3 independent experiments. Ordinary one-way ANOVA with Tukey’s multiple comparisons test. Mean ± SD. **G**. Representative images of a HEK293 cell expressing SPTLC3-Halo (green), SPTLC1-SNAP (magenta), GFP- RhoBTB3 (blue), and Golgi-BFP (grey) and with insets. Yellow arrows denoted SPTLC3 and SPTLC1 co-localized on RhoBTB3-positive Golgi. **H-K**. Representative images of a HEK293 cell expressing CerS2-Halo (magenta; **H**), SMS1-Halo (magenta; **I**), CHPT1-Halo (magenta; **J**), or PEMT-Halo (magenta; **K**), along with GFP-RhoBTB3 (green), and Golgi- BFP (blue) with insets. **L**. Pearson’s correlation coefficient of GFP-RhoBTB3 vs either SPTLC1 (14 cells), CerS2 (12 cells), SMS1 (15 cells), SMS2 (16 cells), PEMT (13 cells), CCTα (13 cells), CEPT (15 cells), CHPT1 (14 cells), CKα (15 cells), or SPK1 (12 cells) from 3 independent experiments. Ordinary one-way ANOVA with Tukey’s multiple comparisons test. Mean ± SD. **M-Q**. GFP-Trap assays demonstrate interactions between GFP-RhoBTB3 and Cers2-Halo (**M**), SMS1-Halo (**N**), CEPT1-Halo (**O**), PEMT-Halo (**P**), CHPT1-Halo (**Q**) in HEK293 cells. Scale bar, 10μm in the whole cell images and 2μm in the insets in (A, C, E, G & H-K).

Interestingly, SPTLCs were reported to be integral ER proteins^45^. Through dissections of the SPTLC3 protein (Fig. S16C), we found that a NT region containing a TM (SPTLC3-1-100) was mainly on the ER (Fig. S16D), while its CT (SPTLC3-174-427) was on the Golgi (Fig. S16E). Consistently, the SPTLC3-174-427 was co-localized with GFP-SHIP164 and Halo-RhoBTB3 on the Golgi (Fig. S16F). These results suggested that SPTLC3 may shuttle between the ER and the Golgi, and RhoBTB3 and SHIP164 may form a complex with SPTLC3 on the Golgi. SPTLC3 forms a heterodimer with SPTLC1 constituting the catalytic core, which generates shorter chain sphingoid bases compared to complexes containing SPTLC2^45^. Indeed, SPTLC1, SPTLC3, and RhoBTB3 were co-localized on the Golgi apparatus (Fig. 6G). In addition, other enzymes of sphingolipid synthesis pathway, including CerS2 and SMS1/2 were co-localized with RhoBTB3 on Golgi vesicles but not the Golgi ribbon (Figs. 6H, I&S18B). Consistently, GFP-trap assays confirmed interactions between RhoBTB3 and CerS2, and SMS1 (Fig. 6M, N).

Importantly, we found that Cholinephosphotransferase 1 (CHPT1) that catalyzes PC biosynthesis from CDP-choline^46^, was partially associated with RhoBTB3 on Golgi vesicles (Fig. 6J). Phosphatidylethanolamine N-methyltransferase (PEMT) catalyzes the three sequential steps of the methylation pathway of PC biosynthesis, and was closely associated with RhoBTB3 on the Golgi (Fig. 6K). The choline/ethanolamine phosphotransferase (CEPT) catalyzes the terminal synthesis of PC in the ER^47^, and appeared to be moderately enriched at the ER-Golgi MCSs (Fig. S17A). Consistently, GFP-trap assays confirmed interactions between RhoBTB3 and PEMT, CHPT1, and CEPT (Fig. 6O-Q). Other enzymes of phospholipid synthesis and metabolism, including CTP:phosphocholine cytidylyltransferase α (CCTα) (Fig. S17C) choline kinase α (CKα) (Fig. S17D) and Sphingosine kinase (SPK1) (Fig. S17E) were not associated with the RhoBTB3-positive Golgi complex.

One of the most noticable feature of these RhoBTB3-labeled Golgi vesicles was motile and frequently contacted EEs, but these two organelles did not fuse. Video analyses showed that majority of EE contacting these RhoBTB3+ Golgi vesicles for ∼60-120s (Fig. S18A-F). In addition, these Golgi vesicles also transiently contacted the ER, with most contacts lasting less than 10s (Fig. S18G-H). Together, our results suggest that RhoBTB3-positive Golgi vesicles are motile phospholipid-synthesizing organelles that may supply phospholipids for EE bud growth.

## Discussion

The roles of bridge lipid transporters in organelle biogenesis have been increasingly appreciated. Yeast Vps13 is required for the prospore formation in yeast ^35, 36^, and strains lacking the protein fail to sporulate^37^. Several groups independently reported that Atg2 is required for the autophagosome biogenesis^23, 28^. Vps13D is suggested to be necessary for peroxisome biogenesis^29^. In this study, we discovered a new type of MCSs between the Golgi and EE, and demonstarted that another bridge lipid transporter SHIP164 was recruited to this Golgi-EE contact. The lipid transfer activity of SHIP164 is required for the growth of EE buds, but not EEs. SHIP164 associated with glycerophospholipids and sphingolipids in HEK293 cells. SHIP164 and RhoBTB3 interact with enzymes for phospholipid synthesis on motile Golgi vesicles, that frequently contact EEs. Together, we propose that lipid transport by SHIP164 from the Golgi to EEs at the contacts promotes the biogenesis of EE buds (Fig. 7). Our findings extend the biological functions of bridge lipid transporters, and identify the biogenesis of EE buds as the functional cellular context of the newly identified bridge lipid transporter SHIP164, which builds up a new link between MCSs and vesicular trafficking.

**Fig. 7.**
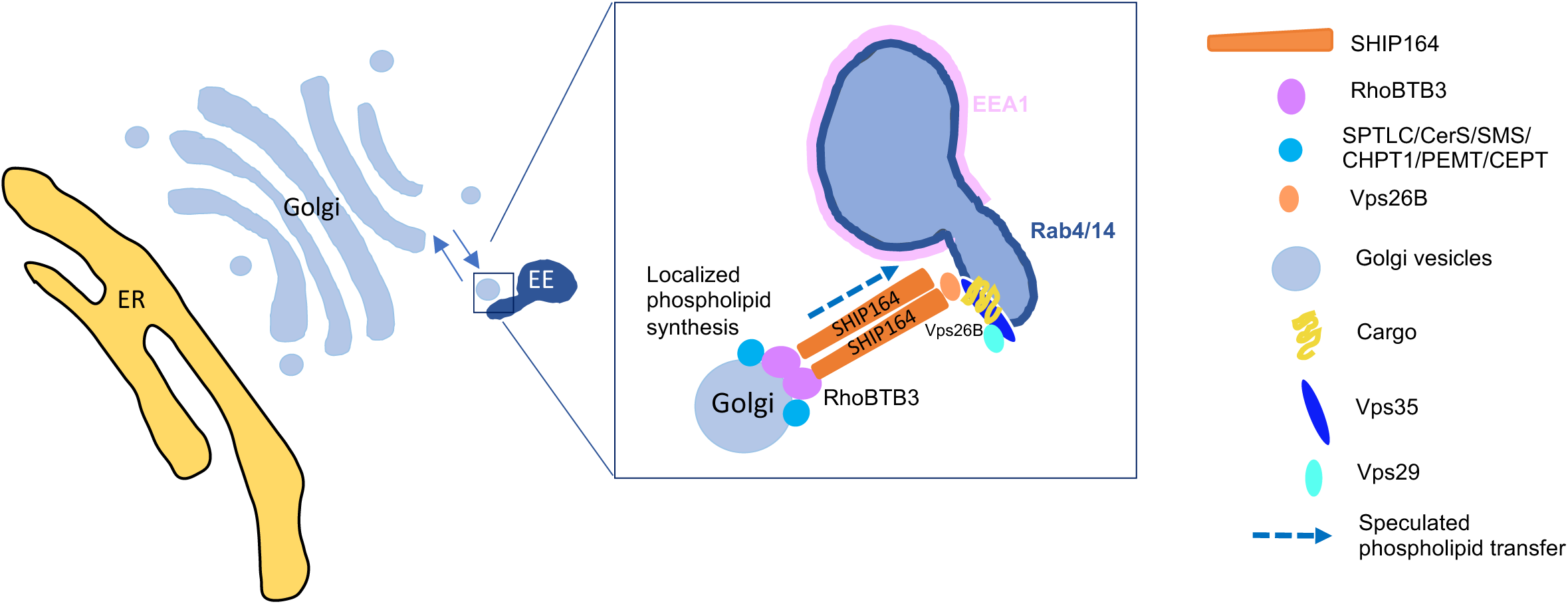
Working model of RhoBTB3-SHIP164-Vps26B at Golgi-EE interface. SHIP164 is recruited to Golgi-EE contacts via two adaptors, an ATPase RhoBTB3 on the Golgi apparatus and a component of retromer Vps26B on EE buds. SHIP164 bridges at the Golgi-EE interface by targeting RhoBTB3 via its NT, meanwhile by recognizing Vps26B via the CT. Functionally, SHIP164 promotes the growth of EE buds, depending on its lipid transfer activity. SHIP164 associates with glycerophospholipids and sphinogolipids, and its depletion reduces LPC/PC and SM level of Rab14-labeled EE membranes, and consequently impairs the SM trafficking to the PM, and affects cell proliferation. SHIP164 and RhoBTB3 interact with enzymes of phospholipid synthesis of on RhoBTB3 positive Golgi vesicles, that frequently contacts EEs, to drive directional lipid transport from the Golgi to EE for EE bud biogenesis. Notably, SHIP164 and RhoBTB3 proteins might be oligomers at the contacts.

It is tempting to speculate that bridge lipid transporters may play a role in the de novo biogenesis of certain organelles or transient membrane structures. It is unclear whether these lipid transporters also play a role in the biogenesis of the organelles that are inherited from mother cells, including the ER, mitochondria, the Golgi, or the plasma membrane, and this fundamental question remains to be answered in the future.

The ER is thought to be the main site for lipid synthesis^43^. Newly synthesized lipids at the ER are then transferred to other organelles through both vesicular and nonvesicular transport pathways. Supporting this notion, many bridge lipid transporters function at ER-associated MCSs. For instance, Vps13A, Vps13C, and Vps13D were suggested to be present at diverse ER MCSs in mammals. Vps13A was shown to localize to ER-mitochondrial, ER-LD, or ER-plasma membrane MCSs^22^ (personal communictions). Vps13C localized to ER-late endosome/lysosome, or ER-LD MCSs^22^. Vps13D localized to ER-mitochondrial, ER-peroxisome MCSs, or mitochondra-LD MCSs upon starvation^16, 27, 29^. Atg2 localized to ER-autophagosome MCSs^23, 28^. The tight associations of these bridge transporters with the ER are in accord with their abilities of binding tens of lipids at once via the extended hydrophobic grooves at the N-terminals^5, 14–16^. However, our results clearly indicated that the localization of SHIP164 was independent of the ER, suggesting that the ER may not be the primary lipid donor organelle for SHIP164. In agreement with this result, our results demonstrated that SHIP164 and its adaptors RhoBTB3 interact with enzymes in the phospholipid synthesis pathways on Golgi vesicles, and these Golgi vesicles were dynamic and contacted EEs in a ‘kiss and run’ manner. These results suggested that these RhoBTB3-positive Golgi vesicles are lipid-synthesizing organelles and supplies phospholipids to EEs. Our findings were mechanistically similar to a previous study by Balla and his colleagues, in which phosphatidylinositol is synthesized in a highly mobile, ER-derived lipid distribution organelle and is delivered to other membranes during multiple contacts^48^.

EEs have been thought to be formed and maintained by the fusion of endocytic vesicles derived from the PM, and then to mature into LEs, which receive TGN-derived vesicles^49^. In this study, our findings identified the existence of a physical contact between the Golgi and EE, and proposed that direct lipid transport from Golgi vesicles to EEs mediated by SHIP164 was required for the growth of EE buds. Our findings were consistent, in part, with a recent study in yeast showing that the formation of EEs depends on a process begun at the Golgi, while endocytic vesicle internalization is not essential for Rab5-mediated endosome formation^50^.

Emerging evidence have shown that bridge lipid transporters cooperate with scramblases to promote membrane expansion^51^. Atg2, proposed to transfer bulk lipids from the ER during autophagosome biogenesis, interacts with two ER-resident scramblases TMEM41B^52, 53^ and VMP1^53^, and with a Golgi-resident scramblase Atg9, that initiate the foramtion of autophagosome^51^. Vps13A was suggested to interact with a PM-resident scramblase XK^54–57^. It is tempting to speculate that SHIP164 may also cooperate with scramblases to mediate the lipid transfer from the Golgi to EEs.

As SHIP164 is highly expressed in the brain, and recent clinical studies indicated that it is involved in Parkinson’s disease^58^ and Myopia, or nearsightedness, a common ocular genetic disease^59^, our findings will provide new mechanistic insights on these synmptoms.

## Materials and methods

### Cell culture, transfection, RNAi

Human embryonic kidney 293T (ATCC) were grown in DMEM (Invitrogen) supplemented with 10% fetal bovine serum (Gibco) and 1% penicillin/streptomycin. The hTERT-immortalized retinal pigment epithelial cell line hTERT RPE-1 cell were grown in DMEM/F12(Gibco) supplemented with 10% fetal bovine serum (Gibco) and 1% penicillin/streptomycin. All of the cell lines used in this study were confirmed free of mycoplasma contamination.

Transfection of plasmids and RNAi oligos was carried out with Lipofectamine 2000 and RNAi MAX, respectively. For transfection, cells were seeded at 4 x 10^5^ cells per well in a six-well dish ∼16 h before transfection. Plasmid transfections were performed in OPTI-MEM (Invitrogen) with 2 μl Lipofectamine 2000 per well for 6 h, followed by trypsinization and replating onto glass-bottom confocal dishes at ∼3.5 x 10^5^ cells per well. Cells were imaged in live-cell medium (DMEM with 10% FBS and 20 mM Hepes with no penicillin or streptomycin) ∼16–24 h after transfection. For all transfection experiments in this study, the following amounts of DNA were used per 3.5 cm well (individually or combined for co-transfection): 1000ng for GFP-SHIP164 and its mutants, 500 ng for Halo-RhoBTB3 and its mutants, OFP-VPS26B; 500 ng for BFP-Golgi, OFP-Rab5A. For siRNA transfections, cells were plated on 3.5 cm dishes at 30–40% density, and 2 μl Lipofectamine RNAimax (Invitrogen) and 50 ng siRNA were used per well. At 48 h after transfection, a second round of transfection was performed with 50 ng siRNAs. Cells were analyzed 24 h after the second transfection for suppression.

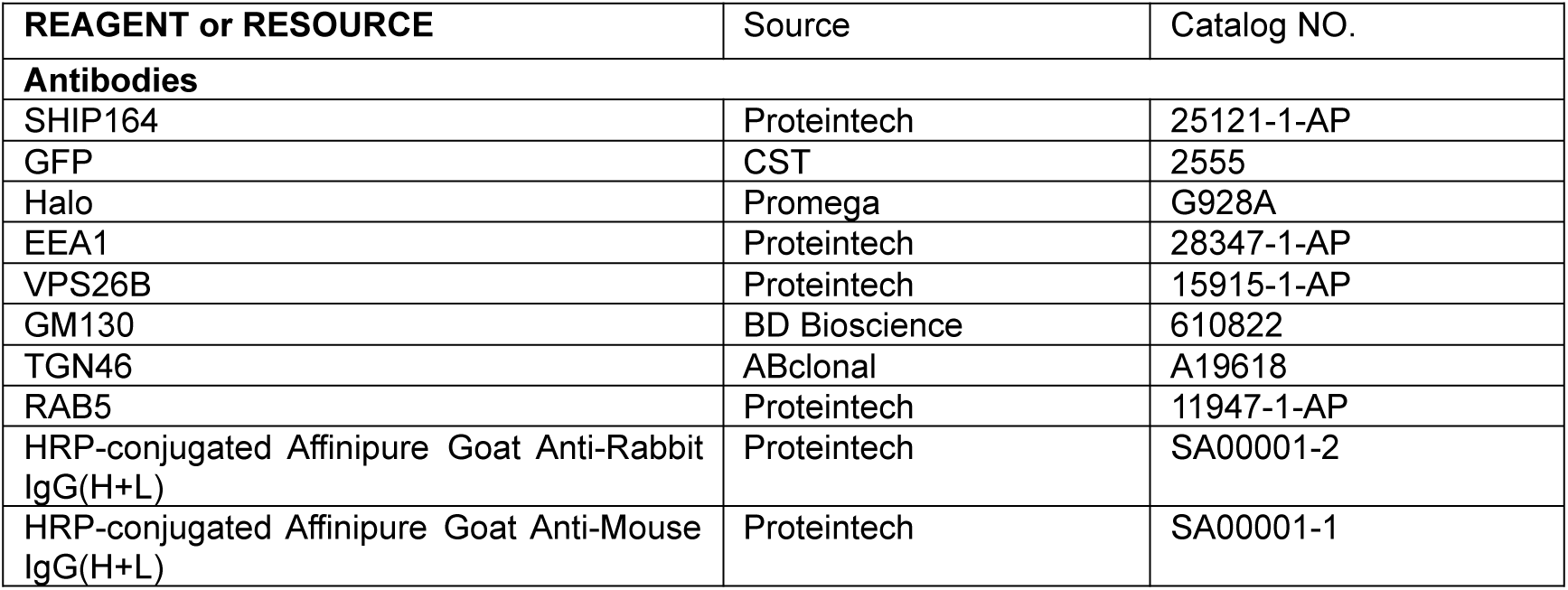
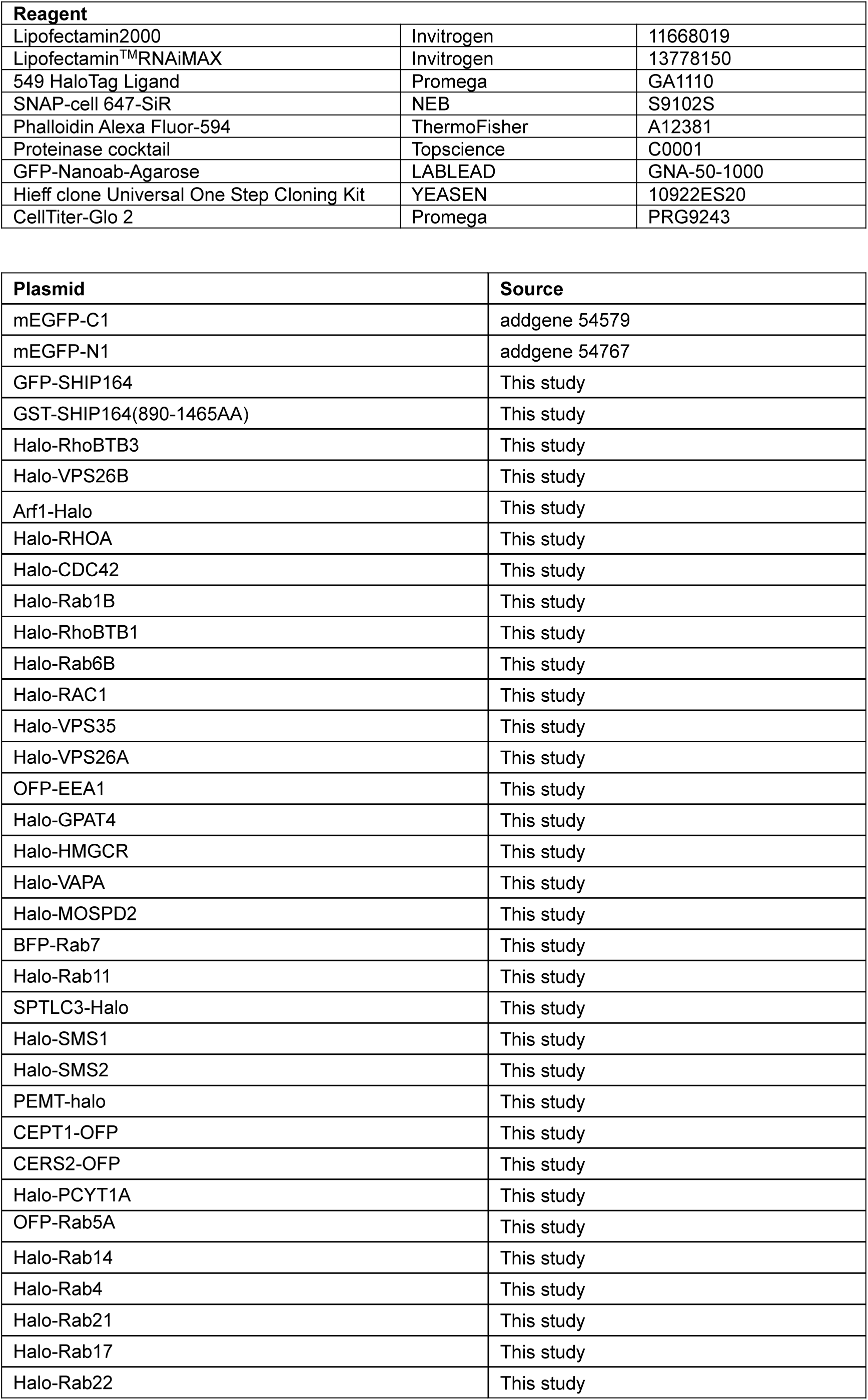
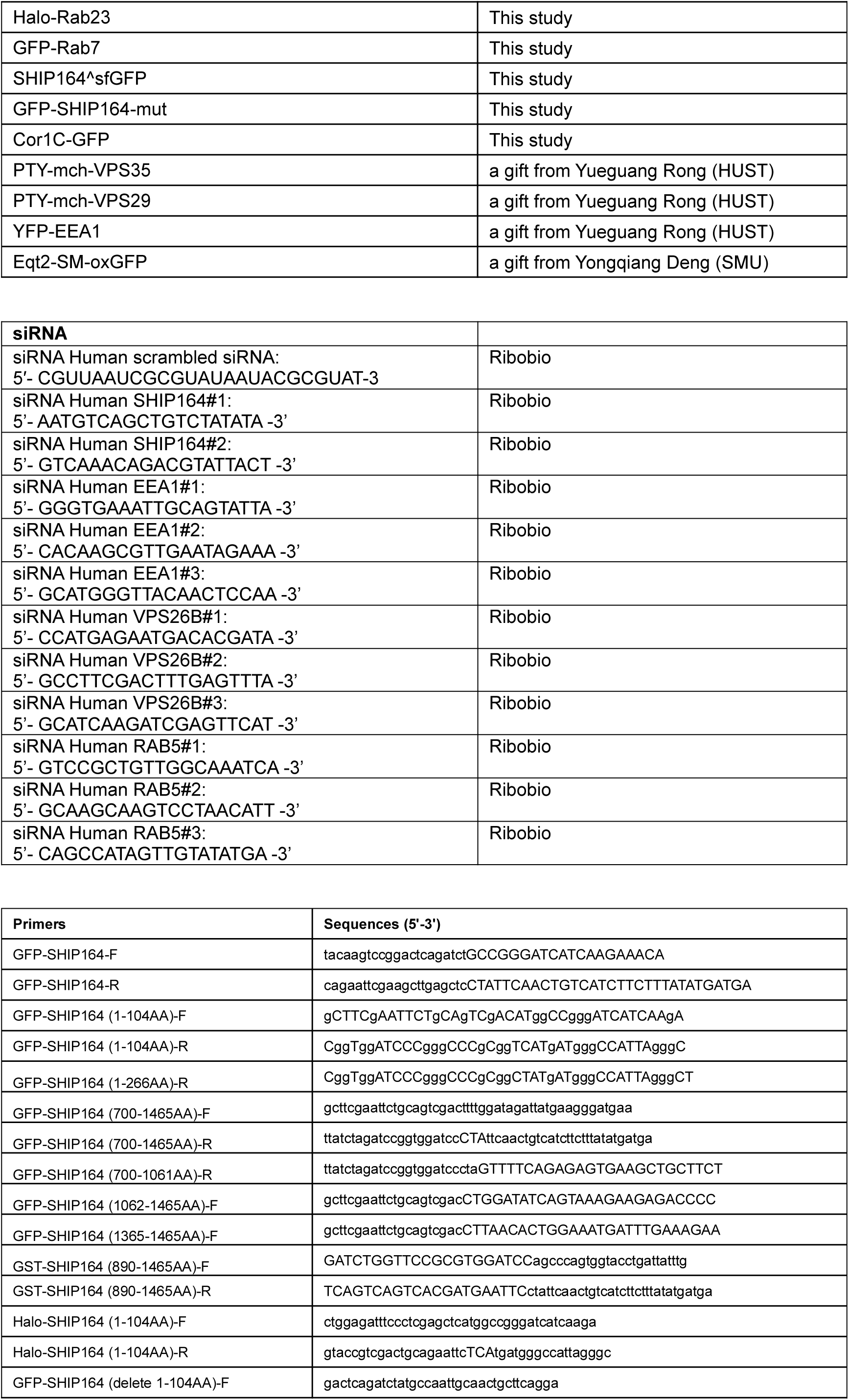
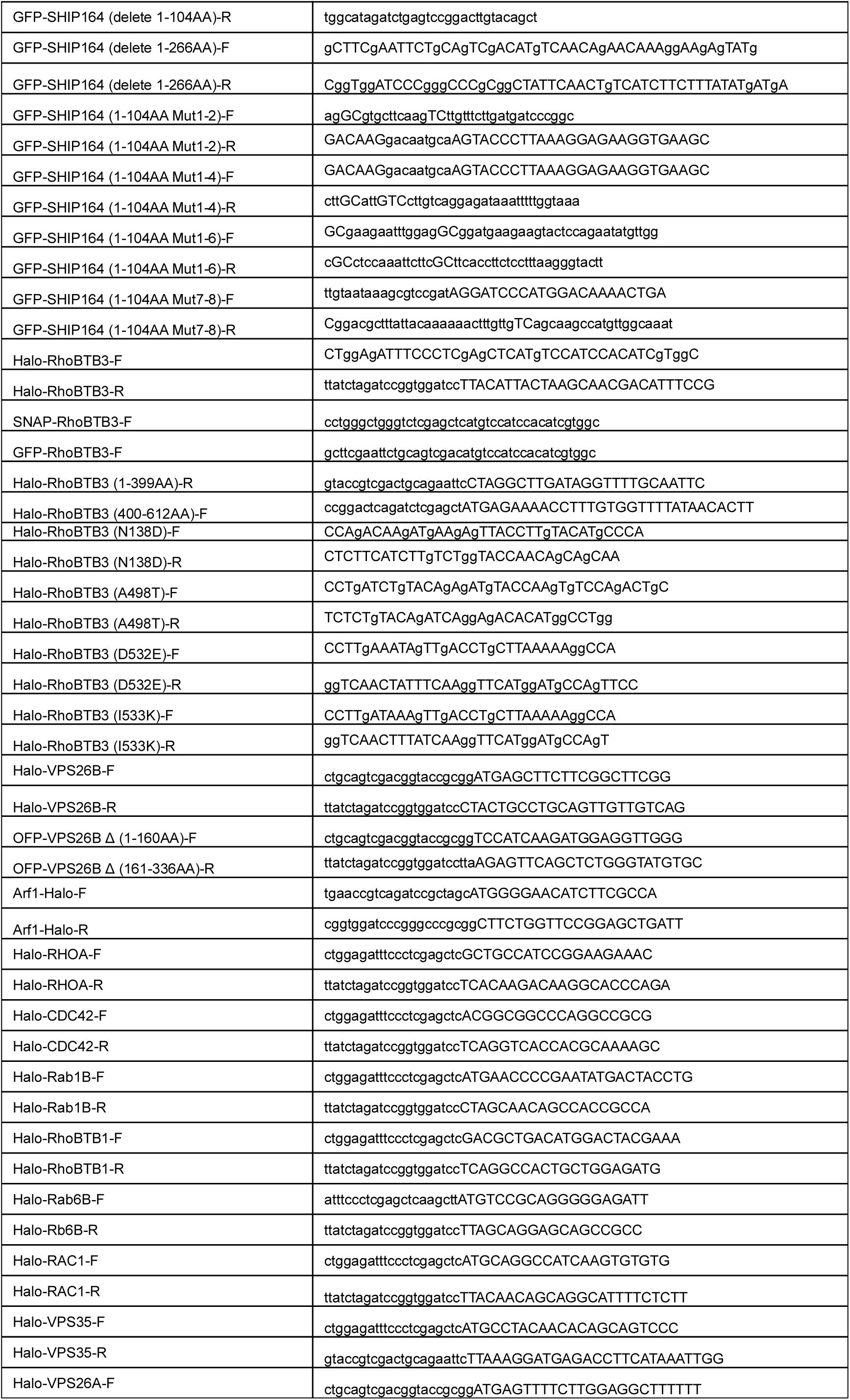
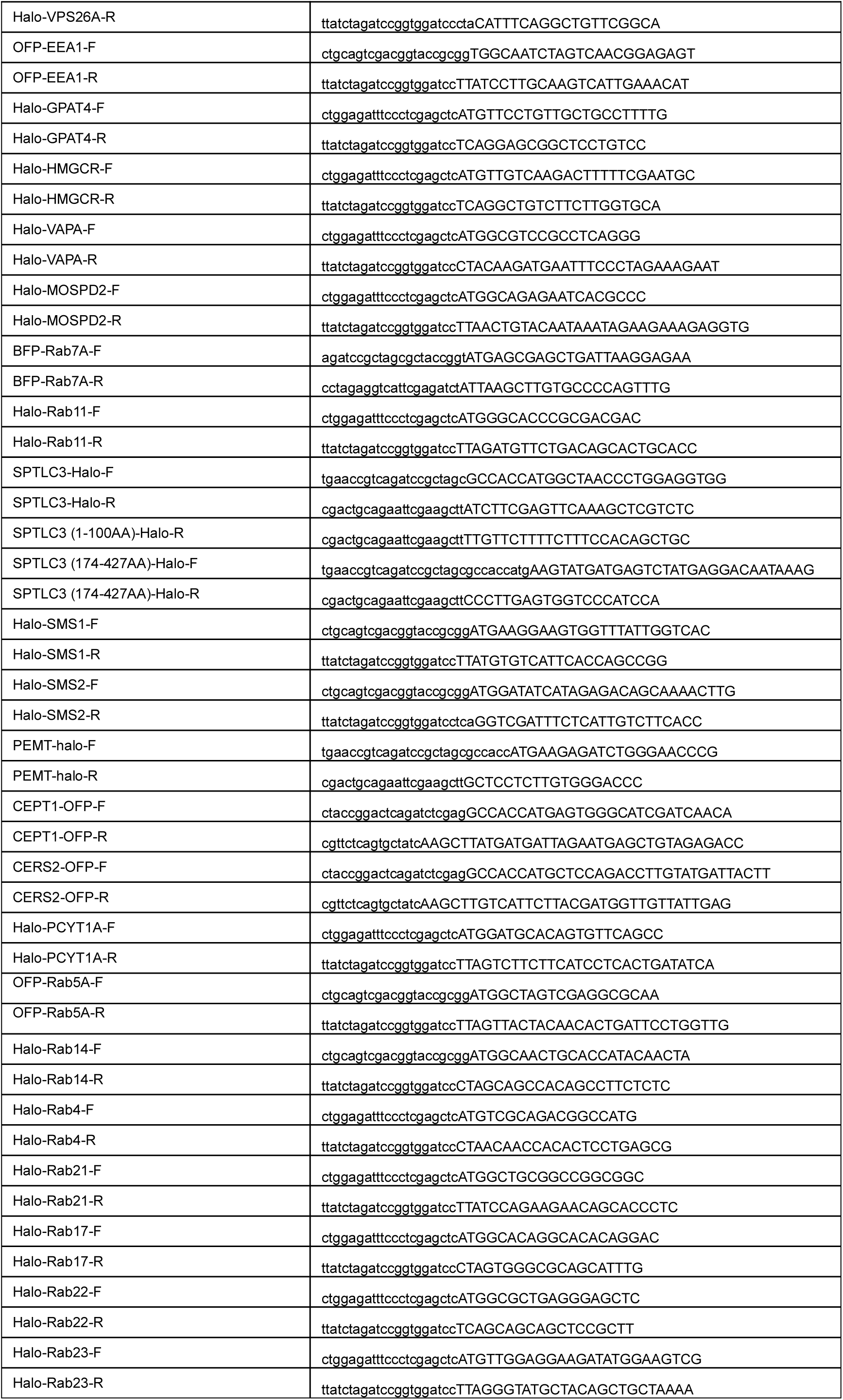
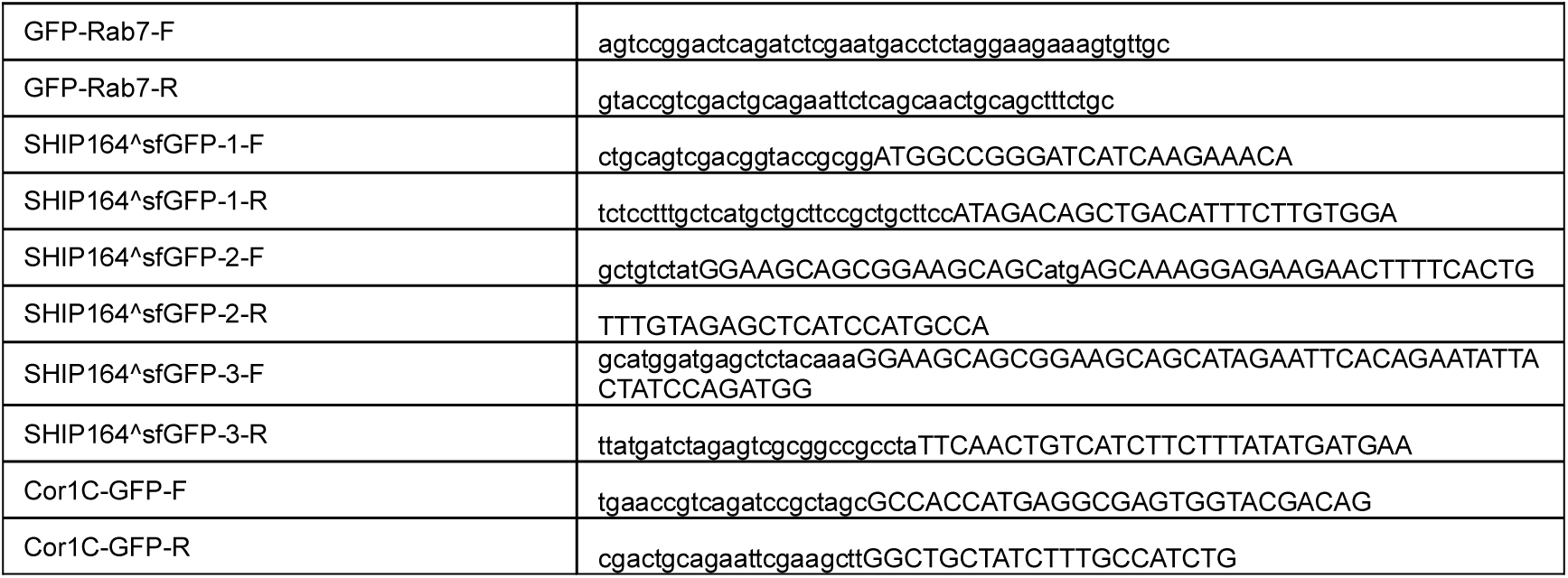

### Duolink PLA Fluorescence Protocol

Cells were fixed with 4% PFA (paraformaldehyde, Sigma) in PBS for 20 min at room temperature. After washing with PBS three times, cells were permeabilized with 0.5% Triton X-100 in PBS for 10 min on ice. Cells were then washed three times with PBS, blocked with Duolink Blocking Solution for 1 h at 37°C. Dilute primary antibodies in the Duolink Antibody Diluent and incubated overnight at 4°C. Wash the slides 2x 5 minutes in 1x Wash Buffer A at room temperature and apply the PLA probe solution for 1 hour at 37 °C. Wash the slides 2x 5 minutes in 1x Wash Buffer A at room temperature and apply the ligation solution for 30min at 37 °C. Wash the slides 2x 5 minutes in 1x Wash Buffer A at room temperature and apply the amplification solution for 100min at 37 °C. Wash the slides 2x 10 minutes in 1x Wash Buffer B at room temperature. Wash the slides in 0.01x Wash Buffer B for 1 minute. samples were mounted on Mounting medium with DAPI.

### GFP-trap assay

GFP trap was used for detection of protein–protein interactions and the GFP-Trap assays were performed according to the manufacturer’s protocol. 5% input was used in GFP traps unless otherwise indicated. Briefly, cells were lysed in Lysis buffer (50 mM Tris-Cl pH 7.5, 150 mM NaCl, 0.5 mM EDTA, 0.5 % Nonidet^TM^ P40 Substitute). After incubation of GFP-Trap beads 4h at 4°C, the beads were pelleted and washed three times with wash buffer (50 mM Tris-HCl pH 7.5, 150 mM NaCl, 0.5 mM EDTA, 1x Proteases Inhibitor cocktail).

### Immunofluorescence staining

Cells were fixed with 4% PFA (paraformaldehyde, Sigma) in PBS for 20 min at room temperature. After washing with PBS three times, cells were permeabilized with 0.5% Triton X-100 in PBS for 10 min on ice. Cells were then washed three times with PBS, blocked with 3% BSA in PBS for 1 h, incubated with primary antibodies in diluted blocking buffer overnight, and washed with PBS three times. Secondary antibodies were applied for 1 h at room temperature. After washing with PBS three times, samples were mounted on Vectashield (H-1000; Vector Laboratories).

### Live imaging by high-resolution confocal microscopy

Cells were grown on 35 mm glass-bottom confocal MatTek dishes, and the dishes were loaded to a laser scanning confocal microscope (LSM900, Zeiss, Germany) equipped with multiple excitation lasers (405 nm, 458 nm, 488 nm, 514 nm, 561 nm and 633 nm) and a spectral fluorescence GaAsP array detector. Cells were imaged with the 63×1.4 NA iPlan-Apochromat 63 x oil objective using the 405 nm laser for BFP, 488 nm for GFP, 561nm for OFP, tagRFP or mCherry and 633nm for Janilia Fluo® 646 HaloTag® Ligand.

### Electron microscopy

HEK293 cells expressing empty vectors or Halo-RhoBTB3, GFP-SHIP164 and OFP- VPS26B were fixed with 2.5% glutaraldehyde in 0.1 M Phosphate buffer, pH 7.4 for 2 h at room temperature. After washing three times with 0.1 M Phosphate buffer, cells were scraped and collected with 0.1 M PBS followed by centrifugation at 1,100 ×g. The pellet was resuspended in PBS (0.1 M), and centrifuged at 1,100 ×g for 10 min. This step was repeated three times. The samples were post-fixed with pre-cold 1% OsO4 in 0.1M PBS for 2-3 hour at 4°C, followed by rinsing with PBS for three times (3 × 20 min). The samples were dehydrated in graded ethanol (50%, 70%, 85%, 90%, 95%, 2x100%) for 15 min in each condition. The penetrations were performed in an order of acetone-epoxy (2:1); acetone-epoxy (1:1); epoxy. Each round of penetrations was performed at 37°C for 12 hours. The samples were embedded in epoxy resin using standard protocols^60^. Sections parallel to cellular monolayers were obtained using a Leica EM UC7 with the thickness of 60-100 nm and examined under HT7800/HT7700.

### Cell viability assay

CellTiter-Glo-2-based cell viability assays were performed according to the manufacturer’s instructions. Briefly, 100 uL detection medium containing 50 uL CellTiter-Glo-2 reagent and 50 uL culture medium was added to one well of 96-well plate containing cells, followed by incubation for 2min on an orbital shaker to break up the cells. After the incubation of the plate at room temperature for 10min to stabilize luminescent signal, luminescence was recorded.

### Mass spectrometry for identification of GST-SHIP164-CT-interacting proteins

GST constructs were transformed into Escherichia coli BL21 (DE3) cells, and cells were incubated at 37°C until the optical density (OD) at 600 nm reached 0.6–0.8, when protein expression was induced with 0.5 mM IPTG, and then cells were cultured at 16°C for another 16 h. Cells were pelleted, resuspended in buffer A (20 mM Tris-Hcl, pH 7.5, 300 mM NaCl, 1mM DTT and 10% glycerol) supplemented with protease inhibitors (Topscience) and lysed via sonication. GST fusion proteins were purified via the GST-tag Protein Purification kit (C600031-0025, Sangon, China). Protein digestion was performed with FASP method. Briefly, the detergent, DTT and IAA in UA buffer was added to block-reduced cysteine. Finally, the protein suspension was digested with 2 µg trypsin (Promega) overnight at 37°C. The peptide was collected by centrifugation at 16,000 g for 15 min. The peptide was desalted with C18 StageTip for further LC-MS analysis. LC-MS/MS experiments were performed on a Q Exactive Plus mass spectrometer that was coupled to an Easy nLC (Thermo Fisher Scientific). Peptide was first loaded to a trap column (100 µm x 20 mm, 5 µm, C18, Dr Maisch GmbH, Ammerbuch, Germany) in buffer A (0.1% formic acid in water). Reverse-phase high- performance liquid chromatography (RP-HPLC) separation was performed using a self-packed column (75 µm x 150 mm; 3 µm ReproSil-Pur C18 beads, 120 Å, Dr Maisch GmbH, Ammerbuch, Germany) at a flow rate of 300 nl/min. The RP-HPLC mobile phase A was 0.1% formic acid in water, and B was 0.1% formic acid in 95% acetonitrile. The gradient was set as following: 2%–4% buffer B from 0 min to 2 min, 4% to 30% buffer B from 2 min to 47 min, 30% to 45% buffer B from 47 min to 52 min, 45% to 90% buffer B from 52 min and to 54 min, and 90% buffer B kept until to 60 min. MS data was acquired using a data-dependent top20 method dynamically choosing the most abundant precursor ions from the survey scan (350– 1800 m/z) for HCD fragmentation. A lock mass of 445.120025 Da was used as internal standard for mass calibration. The full MS scans were acquired at a resolution of 70,000 at m/z 200, and 17,500 at m/z 200 for MS/MS scan. The maximum injection time was set to 50 ms for MS and 50 ms for MS/ MS. Normalized collision energy was 27 and the isolation window was set to 1.6 Th. Dynamic exclusion duration was 60 s. The MS data were analyzed using

MaxQuant software version 1.6.1.0. MS data were searched against the UniProtKB Human norvegicus database (36,080 total entries, downloaded 08/14/2018). Trypsin was seleted as the digestion enzyme. A maximum of two missed cleavage sites and the mass tolerance of 4.5 ppm for precursor ions and 20 ppm for fragment ions were defined for database search. Carbamidomethylation of cysteines was defined as a fixed modification, while acetylation of protein N-terminal, oxidation of Methionine were set as variable modifications for database searching. The database search results were filtered and exported with a <1% false discovery rate (FDR) at peptide-spectrum-matched level, and protein level, respectively.

### Mass spectrometry for identification of GFP-SHIP164-associated lipids in HEK293 cells

The identification of GFP-SHIP164-assocaiting lipids in HEK293 cells was described in our previous study ^21^. Briefly, to extract lipids, 1 ml methyl tert-butyl ether (MTBE) was added to GFP-Trap agarose beads and the samples were shaken for 1 h at room temperature. Next, phase separation was induced by adding 250 µL water, letting it sit for 10 min at room temperature and centrifuging for 15 min at 14,000 g, 4°C. Because of the low density and high hydrophobicity of MTBE, lipids and lipophilic metabolites are mainly extracted to the upper MTBE-rich phase. The lipid was transferred to fresh tubes and dried with nitrogen. Additionally, to ensure data quality for metabolic profiling, quality control (QC) samples were prepared by pooling aliquots from representative samples for all of the analysis samples, and were used for data normalization. QC samples were prepared and analyzed with the same procedure as that for the experiment samples in each batch. Dried extracts were then dissolved in 50% acetonitrile. Each sample was filtered with a disposable 0.22 μm cellulose acetate and transferred into 2 ml HPLC vials and stored at -80°C until analysis. For UHPLC-MS/MS analysis, lipid analysis was performed on Q Exactive orbitrap mass spectrometer (Thermo Fisher Scientific) coupled to a UHPLC system Ultimate 3000 (Thermo Fisher Scientific). Samples were separated using a Hypersil GOLD C18 column (100 x 2.1 mm, 1.9 µm) (Thermo Fisher Scientific). Mobile phase A was prepared by dissolving 0.77 g of ammonium acetate to 400 ml of HPLC-grade water, followed by adding 600 ml of HPLC-grade acetonitrile. Mobile phase B was prepared by mixing 100 ml of acetonitrile with 900 ml isopropanol. The flow rate was set as 0.3 mL/min. The gradient was 30% B for 0.5 min and was linearly increased to 100% in 10.5 min, and then maintained at 100% in 2 min, and then reduced to 30% in 0.1 min, with 4.5 min re-equilibration period employed. Both electrospray ionization (ESI) positive-mode and negative-mode were applied for MS data acquisition. The positive mode of spray voltage was 3.0 kV and the negative mode 2.5 kV. The ESI source conditions were set as follows: heater temperature of 300°C, Sheath Gas Flow rate, 45arb, Aux Gas Flow Rate, 15 arb, Sweep Gas Flow Rate, 1arb, Capillary Temp, 350°C, S-Lens RF Level, 50%. The full MS scans were acquired at a resolution of 70,000 at m/z 200, and 17,500 at m/z 200 for MS/MS scans. The maximum injection time was set to for 50 ms for MS and 50 ms for MS/MS. MS data was acquired using a data-dependent Top10 method dynamically choosing the most abundant precursor ions from the survey scan (200–1500m/z) for HCD fragmentation. Stepped normalized collision energy was set as 15, 25, 35 and the isolation window was set to 1.6 Th. QC samples were prepared by pooling aliquots that were representative of all samples under analysis, and used for data normalization. Blank samples (75% acetonitrile in water) and QC samples were injected every six samples during acquisition.

For data preprocessing and filtering, lipids were identified and quantified using LipidSearch 4.1.30 (Thermo, CA). Mass tolerance of 5 ppm and 10 ppm were applied for precursor and product ions. Retention time shift of 0.25 min was performed in ‘alignment’. M-score and chromatographic areas were used to reduce false positives. The lipids with less than 30% relative standard deviation (RSD) of MS peak area in the QC samples were kept for further data analysis. SIMCAP software (Version 14.0, Umetrics, Sweden) was used for all multivariate data analyses and modeling. Data were mean-centered using Pareto scaling. Models were built on principal component analysis (PCA), orthogonal partial least-square discriminant analysis (PLS-DA) and partial least-square discriminant analysis (OPLS-DA). All the models evaluated were tested for over fitting with methods of permutation tests. The descriptive performance of the models was determined by R2X (cumulative) [perfect model: R2X (cum)=1] and R2Y (cumulative) [perfect model: R2Y (cum)=1] values while their prediction performance was measured by Q2 (cumulative) [perfect model: Q2 (cum)=1] and a permutation test (n=200). The permuted model should not be able to predict classes – R2 and Q2 values at the Y-axis intercept must be lower than those of Q2 and the R2 of the non-permuted model. OPLS-DA allowed the determination of discriminating metabolites using the variable importance on projection (VIP). The VIP score value indicates the contribution of a variable to the discrimination between all the classes of samples. Mathematically, these scores are calculated for each variable as a weighted sum of squares of PLS weights. The mean VIP value is 1, and usually VIP values over 1 are considered as significant. A high score is in agreement with a strong discriminatory ability and thus constitutes a criterion for the selection of biomarkers. The discriminating metabolites were obtained using a statistically significant threshold of variable influence on projection (VIP) values obtained from the OPLS-DA model and two-tailed Student’s t-test (P- value) on the normalized raw data at univariate analysis level. The P-value was calculated by one-way analysis of variance (ANOVA) for multiple groups’ analysis. Metabolites with VIP values greater than 1.0 and P-value less than 0.05 were considered to be statistically significant metabolites. Fold change was calculated as the logarithm of the average mass response (area) ratio between two arbitrary classes. On the other side, the identified differential metabolites were used to perform cluster analyses with R package.

### Statistical analysis

All statistical analyses and p-value determinations were performed in GraphPad Prism6. All the error bars represent Mean ± SD. To determine p-values, ordinary one-way ANOVA with Tukey’s multiple comparisons test were performed among multiple groups and a two-tailed unpaired student t-test was performed between two groups.

## Acknowledgements

We thank Anbing Shi (HUST, China) for critical discussions. We thank Yongqiang Deng (Southern Medical University, China) for discussion and reagents. We thank Hao Chen (SUSTech, China) for assistance in lipidomics. We thank the Mass Spectrometry Core Facility (Mr. Cookson K. C. Chiu), the Bio-imaging Core Facility (Dr. Zhenglong Sun and Ms. Mei Yu), and the Sequencing Core Facility (Dr. Miao Cui) of Shenzhen Bay Laboratory for providing technical supports. We thank Qing Tian and Linfang Yang (HUST, China) and Ping Liu (the Optical Bioimaging Core Facility of WNLO-HUST) for imaging assistance.

## Author contributions

J.W. and W.J. conceived the project and designed the experiments. J.W. performed the experiments J.W., Q.C., W.C., L.D., and W.J. analyzed and interpreted the data. W.J. prepared the manuscript with inputs and approval from all authors.

## Data and materials availability

All the data and relevant materials, including reagents and primers, that supports the findings of this study are available from the corresponding author upon reasonable request.

---

**Fig. S1.**
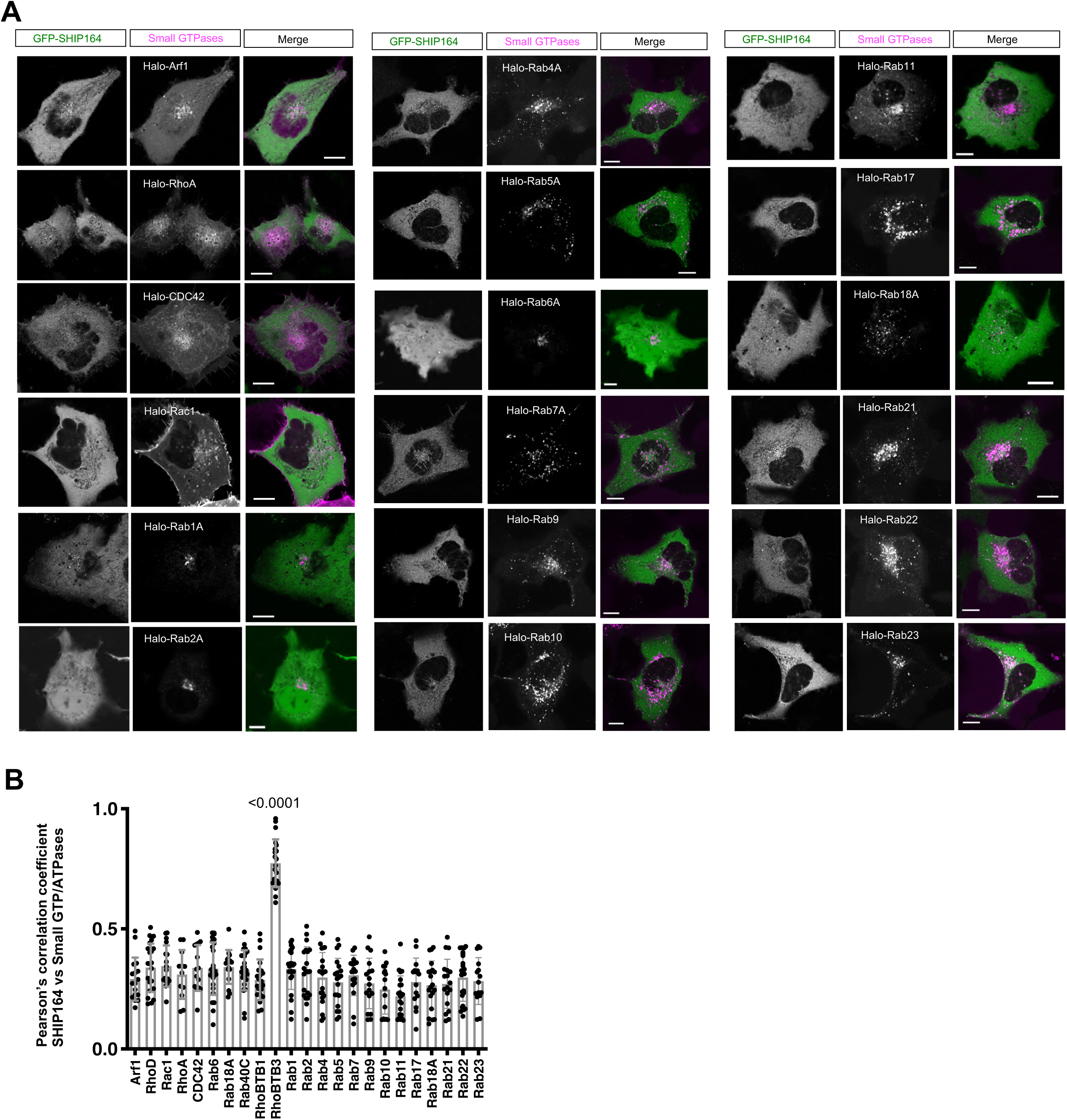
Screening of SHIP164 adaptors using small GTPase library. **A.** Representative images of live HEK293 cells expressing GFP-SHIP164 (green) and small GTPases (magenta). **B.** Pearson’s correlation coefficient of GFP-SHIP164 vs Arf1(17 cells); RhoA (13 cells); CDC42 (16 cells); Rac1 (18 cells); Rab1A (26 cells); Rab2A (22 cells); Rab4A (23 cells); Rab5A (21 cells); Rab6A (26 cells); Rab7A (19 cells); Rab9 (18 cells); Rab10 (17 cells); Rab11 (23 cells); Rab17 (16 cells); Rab18A (21 cells); Rab40c (27 cells); RhoBTB1 (23 cells); RhoBTB3 (21 cells); Rab21 (19 cells); Rab22 (26 cells); and Rab23 (16 cells) in more than 3 independent experiments. Ordinary one-way ANOVA with Tukey’s multiple comparisons test. Mean ± SD. Scale bar, 10μm in the whole cell images and 2μm in the insets in (A).

**Fig. S2.**
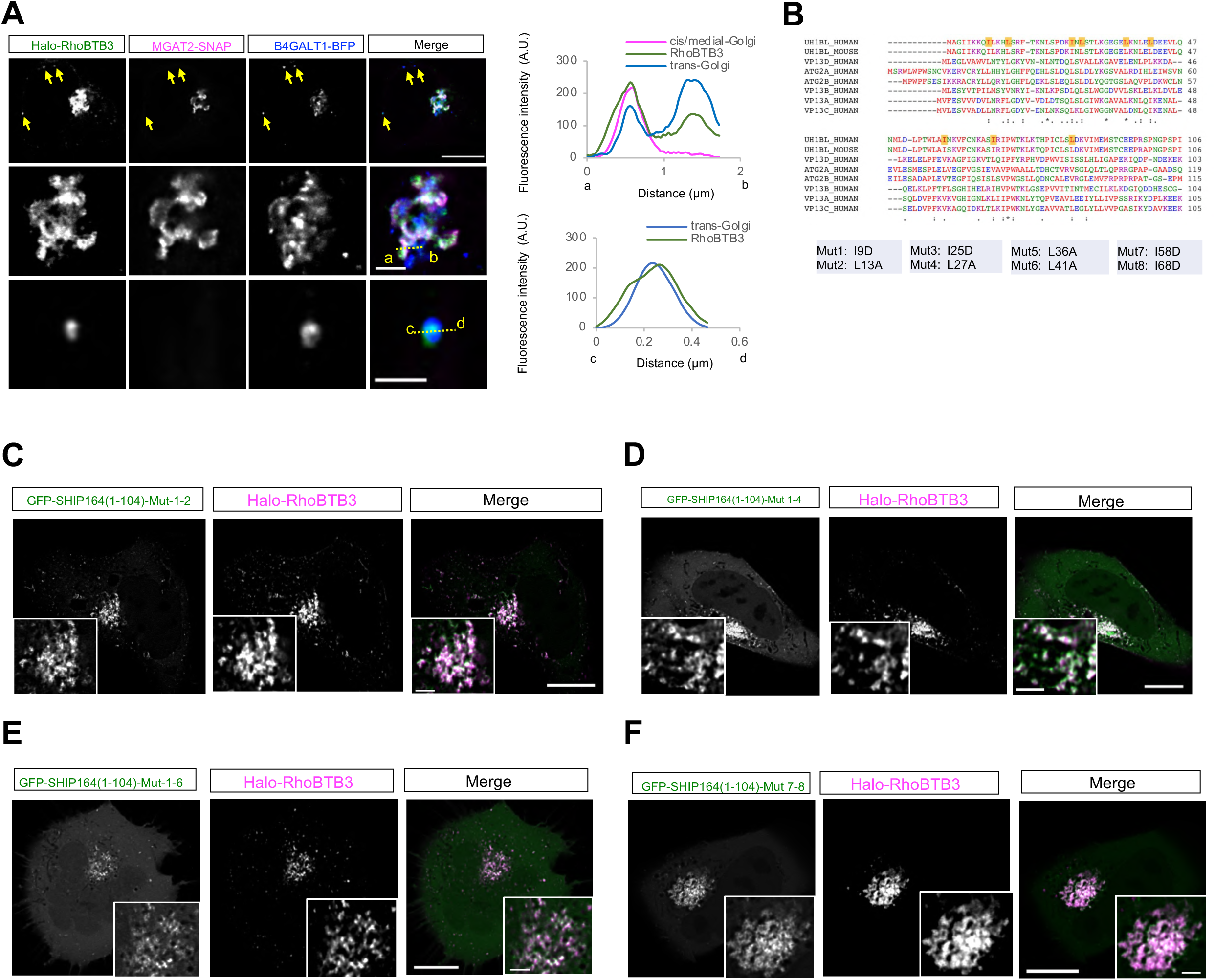
Supplemental results to Figure 1. **A.** Representative images of a live HEK293 cell expressing Halo-RhoBTB3 (green), MGAT2-SNAP (magenta), and B4GALT1-BFP (blue) with line-scan analyses on the right. **B.** Sequence alignments of the NT of Vps13 domain containing proteins. 8 conserved residues were shown at the bottom. **C-F**. Representative images of a HEK293 cells expressing either GFP-SHIP164-1-104- Mut1-2 (green; **C**), GFP-SHIP164-1-104-Mut1-4 (green; **D**), GFP-SHIP164-1-104-Mut1-6 (green; **E**), or GFP-SHIP164-1-104-Mut7-8 (green; **F**) and Halo-RhoBTB3 (magenta) with insets. Scale bar, 10μm in the whole cell images and 2μm in the insets in (A & C-F).

**Fig. S3.**
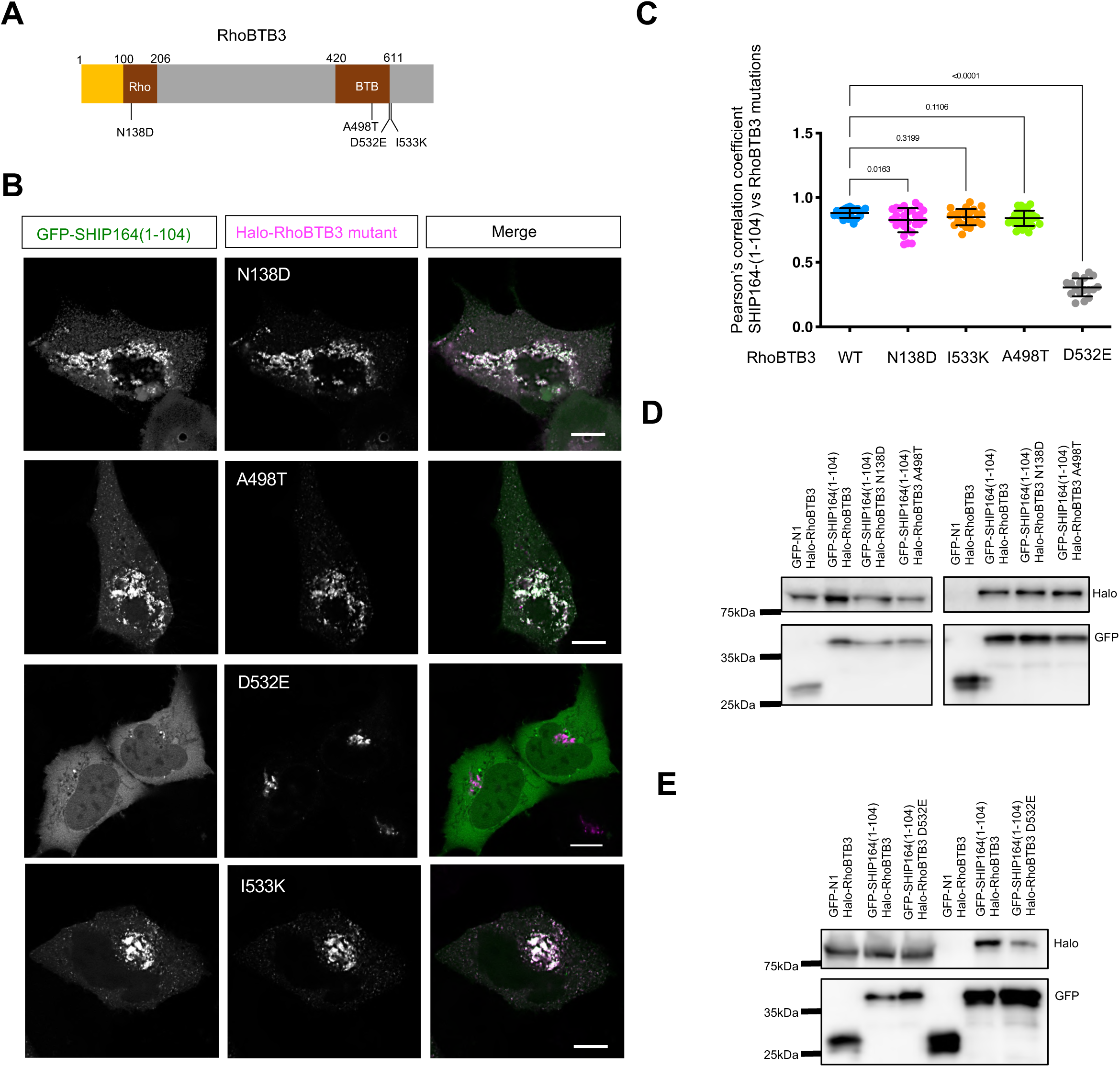
The D532 residue is critical for the RhoBTB3-SHIP164 interaction. **A.** Domain organization of RhoBTB3 with point mutations. **B.** Representative images of live HEK293 cells expressing Halo-RhoBTB3 mutants, N138D/A498T/D532E/I533K (magenta), and GFP-SHIP164-1-104 (green). **C.** Pearson’s correlation coefficient of GFP-SHIP164-1-104 vs Halo-RhoBTB3 mutants: WT Halo-RhoBTB3 (22 cells); N138D (27 cells); I533K (23 cells); A488T (29 cells); or D532E (17 cells) in > 3 independent experiments. Ordinary one-way ANOVA with Tukey’s multiple comparisons test. Mean ± SD. **D-E**. GFP-Trap assays demonstrate interactions between GFP-SHIP164-1-104 and Halo- RhoBTB3 mutants in HEK293 cells. Scale bar, 10μm in the whole cell images in (B).

**Fig. S4.**
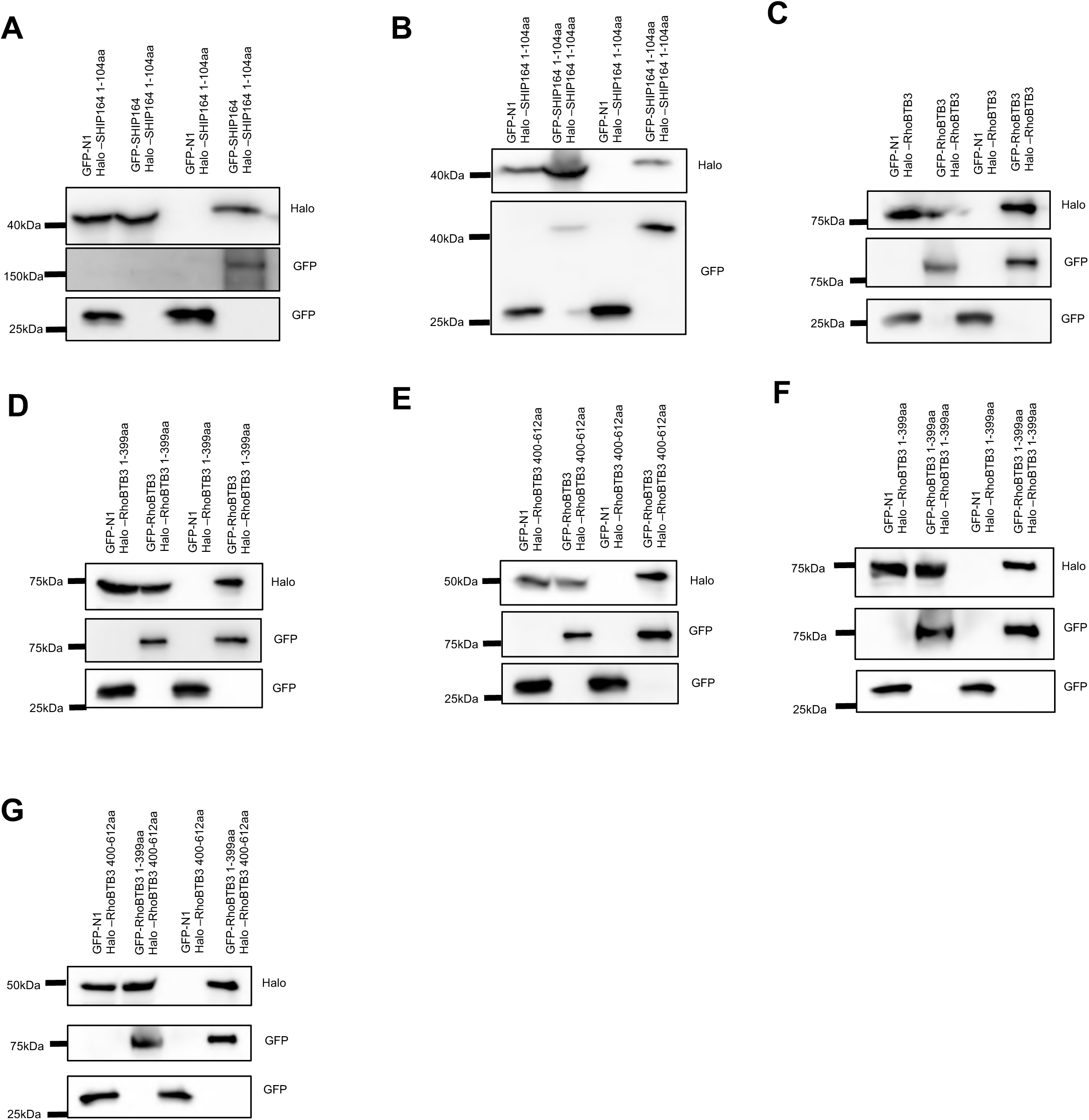
The self-interactions of SHIP164 or RhoBTB3. **A-B**. GFP-Trap assays demonstrated interactions between either GFP-SHIP164 (**A**), or GFP-SHIP164-1-104 (**B**) and Halo-SHIP164-1-104 in HEK293 cells. **C-E**. GFP-Trap assays showed interactions between GFP-RhoBTB3 and either Halo- RhoBTB3 (**C**), Halo-RhoBTB3-1-399 (**D**), or Halo-RhoBTB3-400-612 (**E**) in HEK293 cells. **F-G**. GFP-Trap assays showed interactions between GFP-RhoBTB3-1-399 and either Halo-RhoBTB3-1-399 (**F**), or Halo-RhoBTB3-400-612 (**G**) in HEK293 cells.

**Fig. S5.**
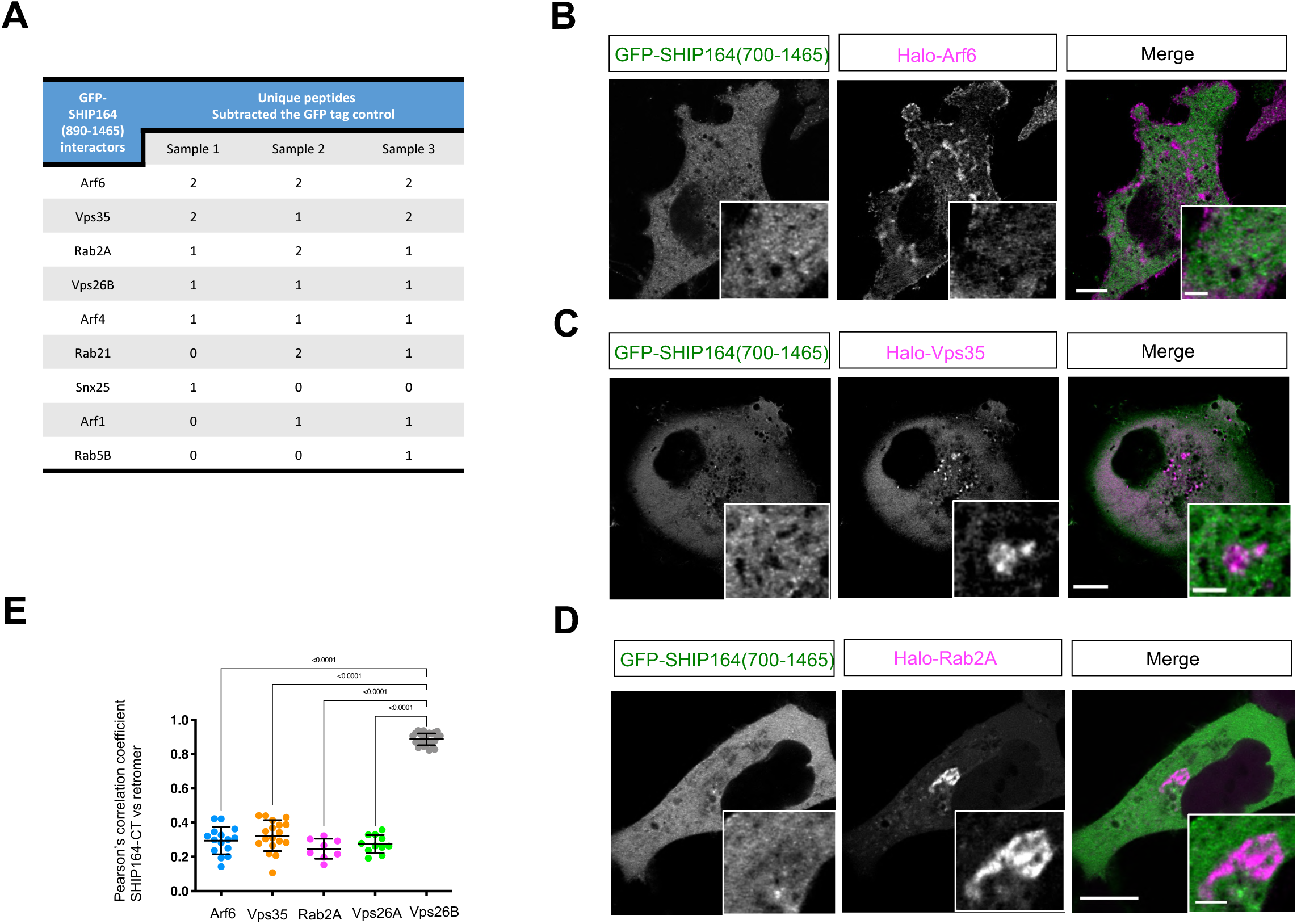
Supplemental results to Figure 2. **A**. A list of protein candidates that functionally related to the endosome-Golgi trafficking, after removal of proteins pull-downed by GST tag alone. **B-D**. Representative images of HEK293 cells expressing GFP-SHIP164-CT (green) and either Halo-Arf6 (magenta; **B**), Halo-Vps35(magenta; **C**), or Halo-Rab2A (magenta; **D**) with insets. **E**. Pearson’s correlation coefficient of GFP-SHIP164-CT vs either Halo-Arf6 (19 cells), Halo-Vps35 (16 cells), Halo-Rab2A (9 cells), Halo-Vps26A (12 cells), or Halo-Vps26B (28 cells) in 3 independent experiments. Ordinary one-way ANOVA with Tukey’s multiple comparisons test. Mean ± SD. Scale bar, 10μm in the whole cell images and 2μm in the insets in (B-D).

**Fig. S6.**
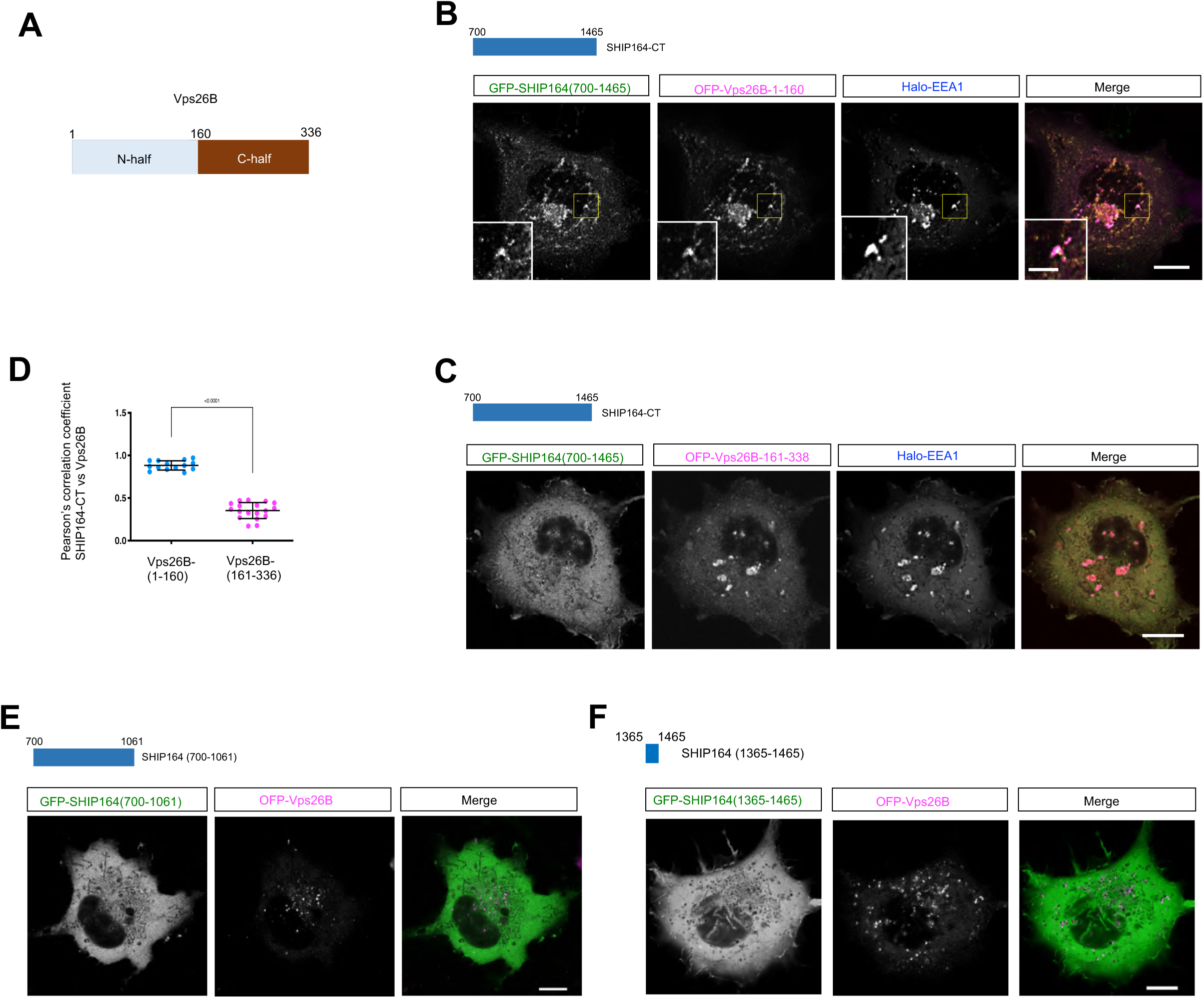
The SHIP164-Vps26B interaction occurs between SHIP164-CT and Vps26B- NT. **A**. Dissection of Vps26B protein. **B-C**. Representative images of HEK293 cells expressing either OFP-Vps26B-1-160 (magenta; B), or OFP-Vps26B-161-338 (magenta; C) and GFP-SHIP164-CT (green) with insets. **D**. Pearson’s correlation coefficient of GFP-SHIP164-CT vs either OFP-Vps26B-1-160 (14 cells), or OFP-Vps26B-161-338 (18 cells) in more than 3 independent experiments. Ordinary one-way ANOVA with Tukey’s multiple comparisons test. Mean ± SD. **E-F**. Representative images of HEK293 cells expressing either GFP-SHIP164-700-1061 (green; **E**), or GFP-SHIP164-1365-1465 (green; **F**) with insets. Scale bar, 10μm in the whole cell images and 2μm in the insets in (B, C, E, & F).

**Fig. S7.**
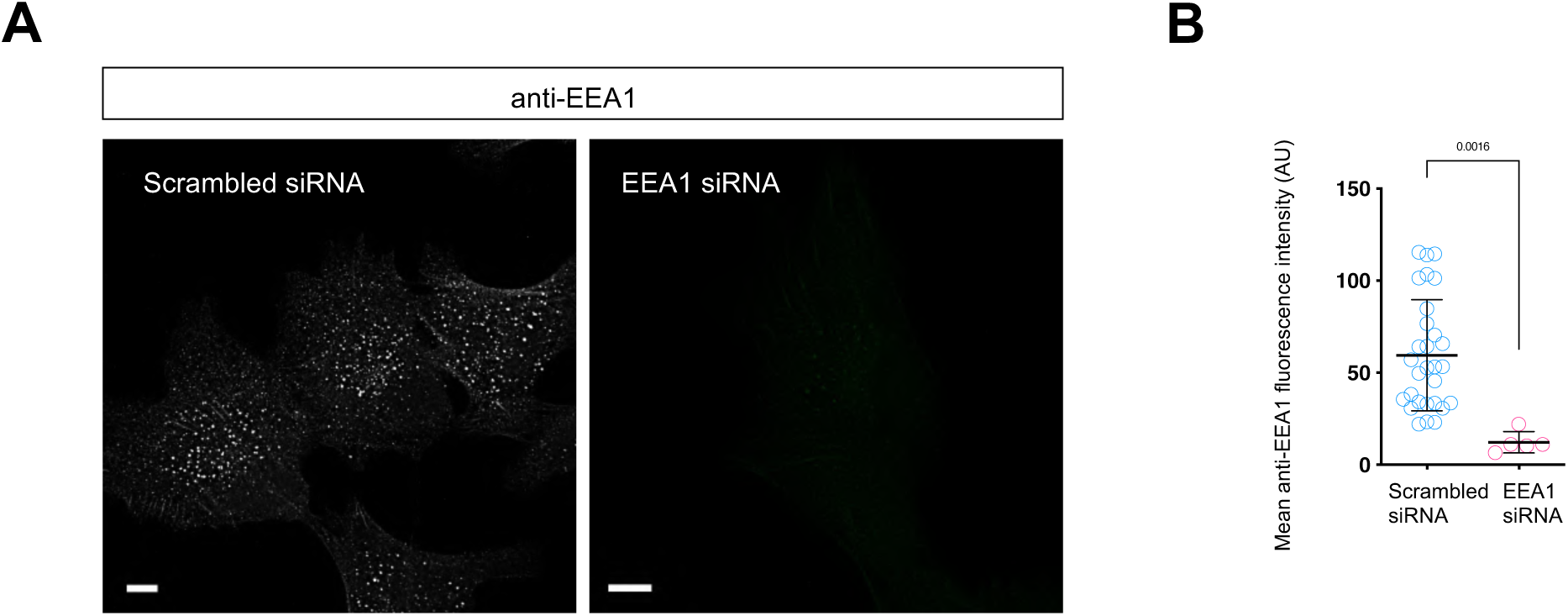
Supplemental results to Figure 3. **A**. Representative images of fixed RPE1 cells stained with antibodies against endogenous EEA1 upon scrambled or EEA1 siRNA. B. Mean fluorescence of anti-EEA1 in scrambled (29 cells) or EEA1 siRNA-treated cells (5 cells). Two-tailed unpaired student t-test. Mean ± SD. Scale bar, 10μm.

**Fig. S8.**
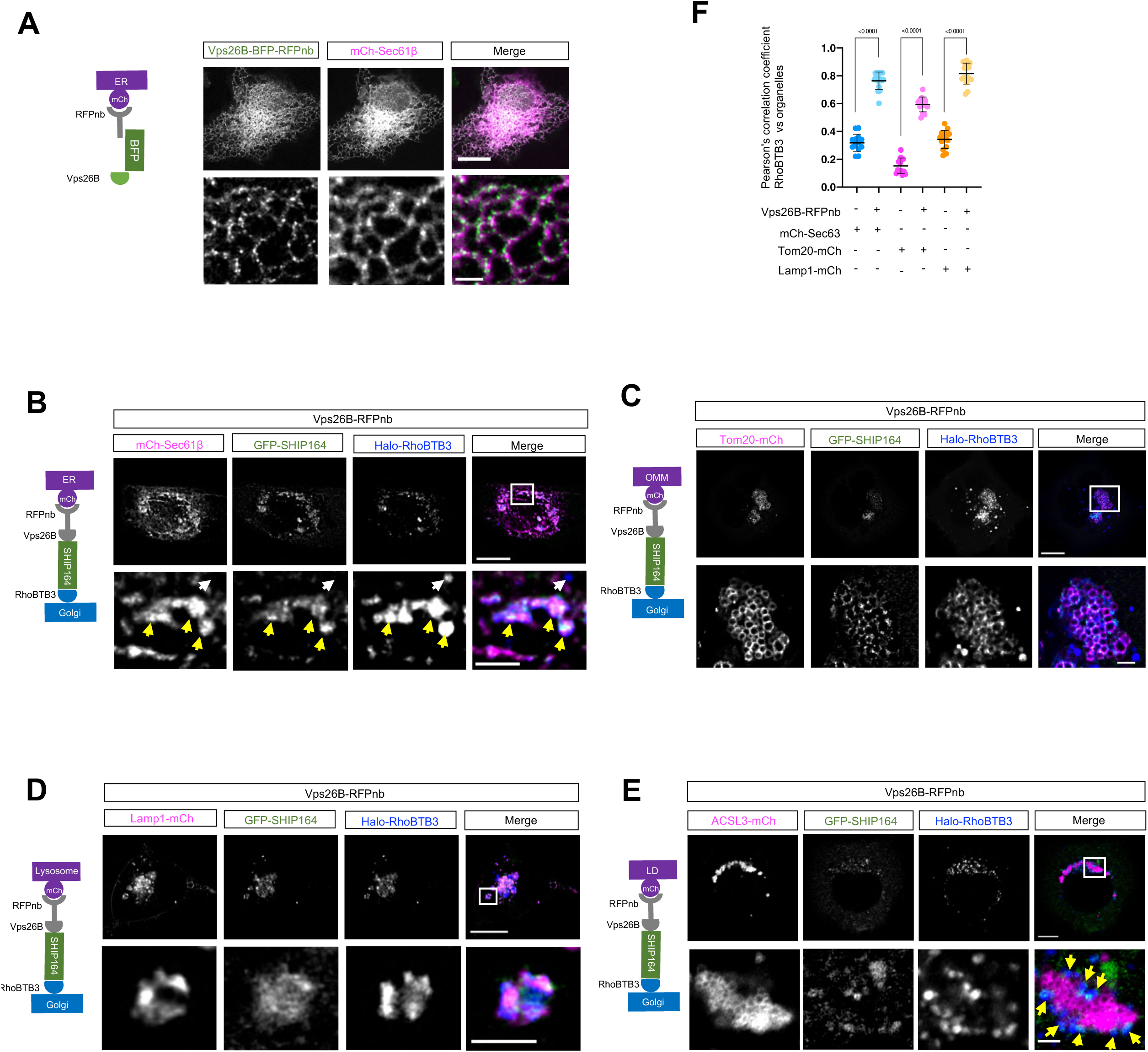
The SHIP164-RhoBTB3-Vps26B complex acts as a tether in cells. **A.** Left: schematic cartoon of RFPnb-mediated recruitment of Vps26B-BFP-RFPnb to mCh-Sec61β-labeled ER membranes. Right: representative images of a HEK293 cell expressing Vps26B-BFP-RFPnb (green) and mCh-Sec61β with insets on the bottom. **B.** Left: schematic cartoon of RFPnb-mediated recruitment of Vps26B-RFPnb to mCh- Sec61β-labeled ER membranes in presence of GFP-SHIP164 and Halo-RhoBTB3. Right: representative images of a HEK293 cell expressing Vps26B-BFP-RFPnb, mCh-Sec61β (magenta), GFP-SHIP164 (green) and Halo-RhoBTB3 (blue) with insets. **C.** Left: schematic cartoon of RFPnb-mediated recruitment of Vps26B-RFPnb to Tom20- mCh-labeled OMM in presence of GFP-SHIP164 and Halo-RhoBTB3. Right: representative images of a HEK293 cell expressing Vps26B-BFP-RFPnb, Tom20-mCh (magenta), GFP-SHIP164 (green) and Halo-RhoBTB3 (blue) with insets. **D.** Left: schematic cartoon of RFPnb-mediated recruitment of Vps26B-RFPnb to Lamp1- mCh-labeled OMM in presence of GFP-SHIP164 and Halo-RhoBTB3. Right: representative images of a HEK293 cell expressing Vps26B-BFP-RFPnb, Lamp1-mCh (magenta), GFP-SHIP164 (green) and Halo-RhoBTB3 (blue) with insets. **E.** Left: schematic cartoon of RFPnb-mediated recruitment of Vps26B-RFPnb to ACSL3- mCh-labeled OMM in presence of GFP-SHIP164 and Halo-RhoBTB3. Right: representative images of an oleic acid-stimulated HEK293 cell expressing Vps26B-BFP- RFPnb, Tom20-mCh (magenta), GFP-SHIP164 (green) and Halo-RhoBTB3 (blue) with insets. **F.** Pearson’s correlation coefficient of Halo-RhoBTB3 vs either mCh-Sec61β (14 cells) or Tom20-mCh (15 cells) in presence of Vps26B-BFP-RFPnb. As controls, Pearson’s correlation coefficient of Halo-RhoBTB3 vs either mCh-Sec61β (13 cells) or Tom20-mCh (15 cells) in absence of Vps26B-BFP-RFPnb. More than 3 independent experiments. Ordinary one-way ANOVA with Tukey’s multiple comparisons test. Mean ± SD. Scale bar, 10μm in the whole cell images and 2μm in the insets in (A-E).

**Fig. S9.**
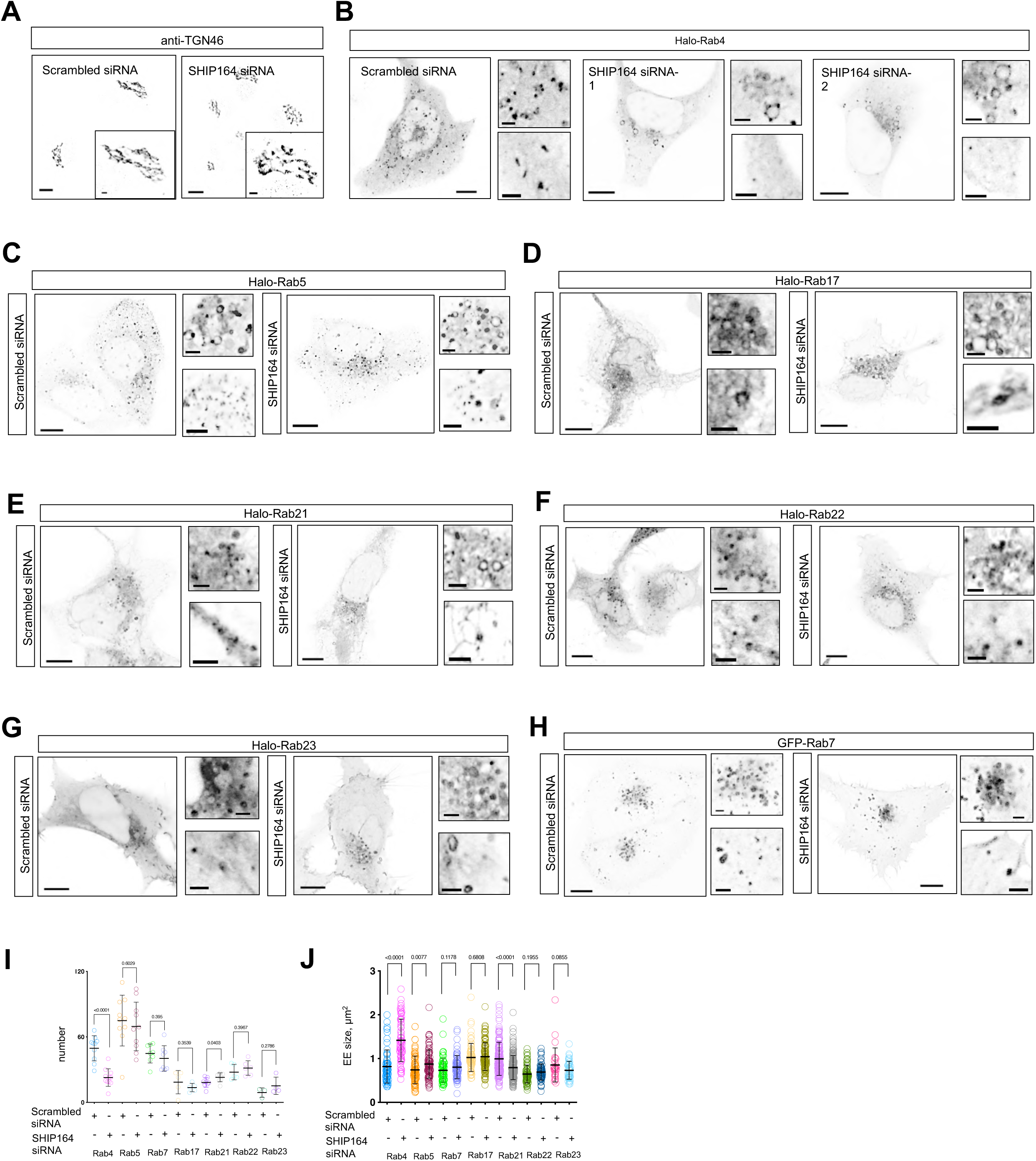
The effects of SHIP164 depletion on the size and number of EE subpopulations. **A**. Representative images of fixed RPE1 cells stained with antibodies against endogenous TGN46 upon scrambled or SHIP164 siRNA. **B-H**. Representative images of HEK293 cells expressing multiple markers for endosomal subpopulations, including Halo-Rab4 (**A**), Halo-Rab5 (**B**), Halo-Rab17 (**C**), Halo-Rab21 (**D**), Halo-Rab22 (**E**), Halo-Rab23 (**F**), or GFP-Rab7 (**G**), upon scrambled or SHIP164 siRNA with two insets. **I, J**. The number (**H**) or size (**I**) of EEs per cell in scrambled or SHIP164 siRNA-treated cells. More than 20 cells from 3 independent assays were quantified for each condition. Ordinary one-way ANOVA with Tukey’s multiple comparisons test. Mean ± SD. Scale bar, 10μm in the whole cell images and 2μm in the insets in (A-H).

**Fig. S10.**
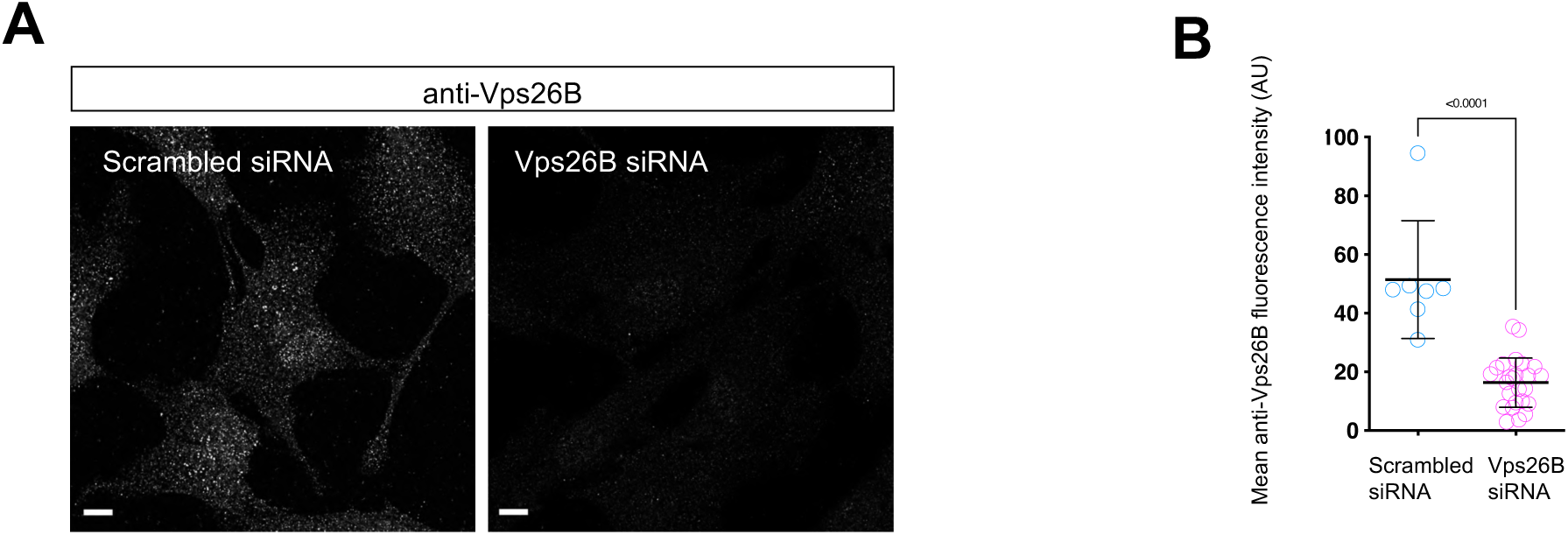
Supplemental results to Figure 4. **A.** Representative images of fixed RPE1 cells stained with antibodies against endogenous Vps26B upon scrambled or Vps26B siRNA. **B.** Mean fluorescence intensity of anti-Vps26B in scrambled (7 cells) or Vps26B depleted cells (25 cells). Scale bar, 10μm in A.

**Fig. S11.**
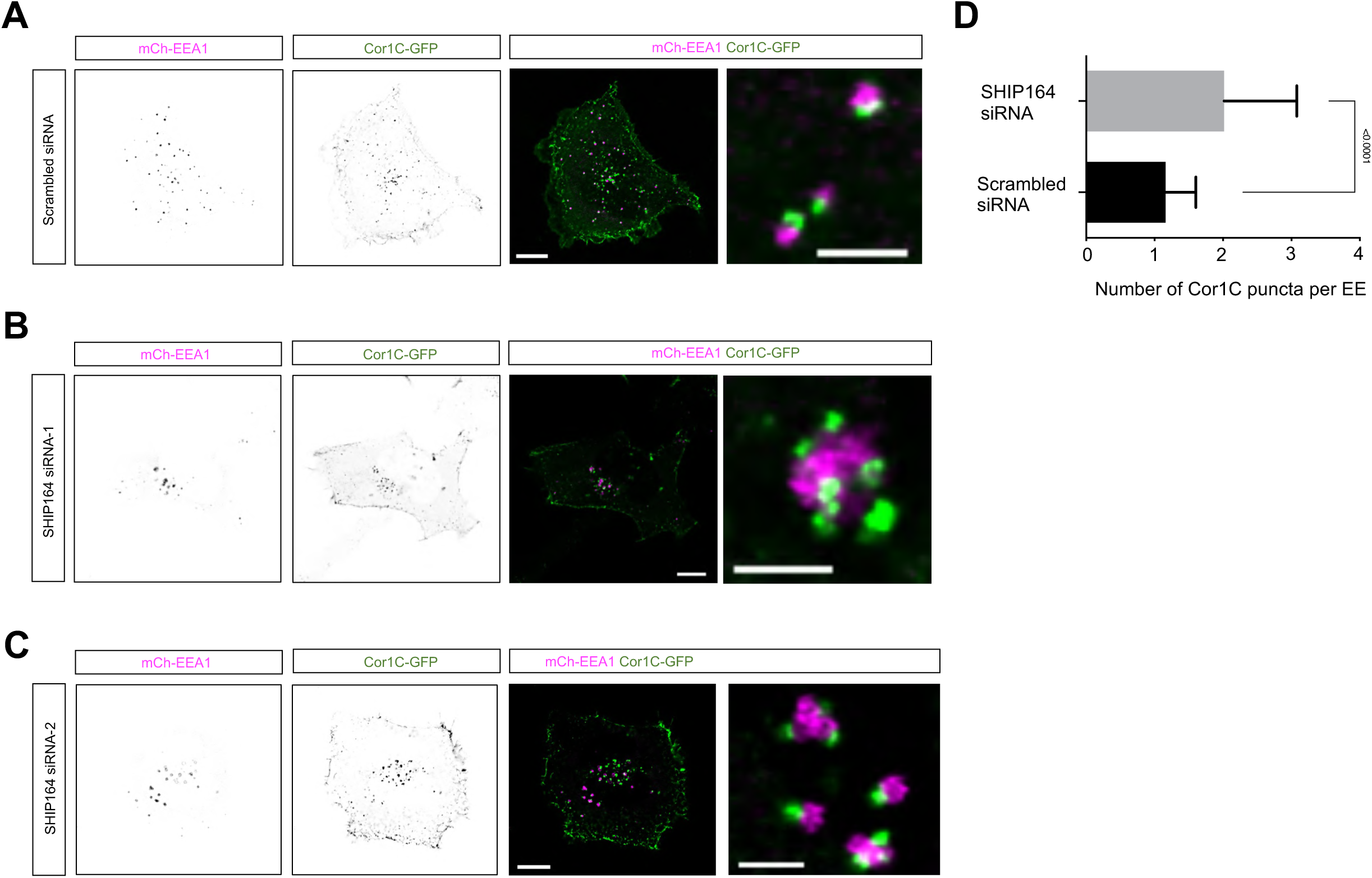
The effect of SHIP164 depletion on Cor1C foci on EE. **A-C**. Representative images of HEK293 cells expressing mCh-EEA1 (magenta) and Cor1C-GFP upon scrambled or SHIP164 siRNA with two insets. **D**. The number of Cor1C puncta per EE in scrambled (14 cells) or SHIP164 siRNA(15 cells)-treated cells from 3 independent assays. Two-tailed unpaired student t-test. Mean ± SD. Scale bar, 10μm in the whole cell images and 2μm in the insets in (A-C).

**Fig. S12.**
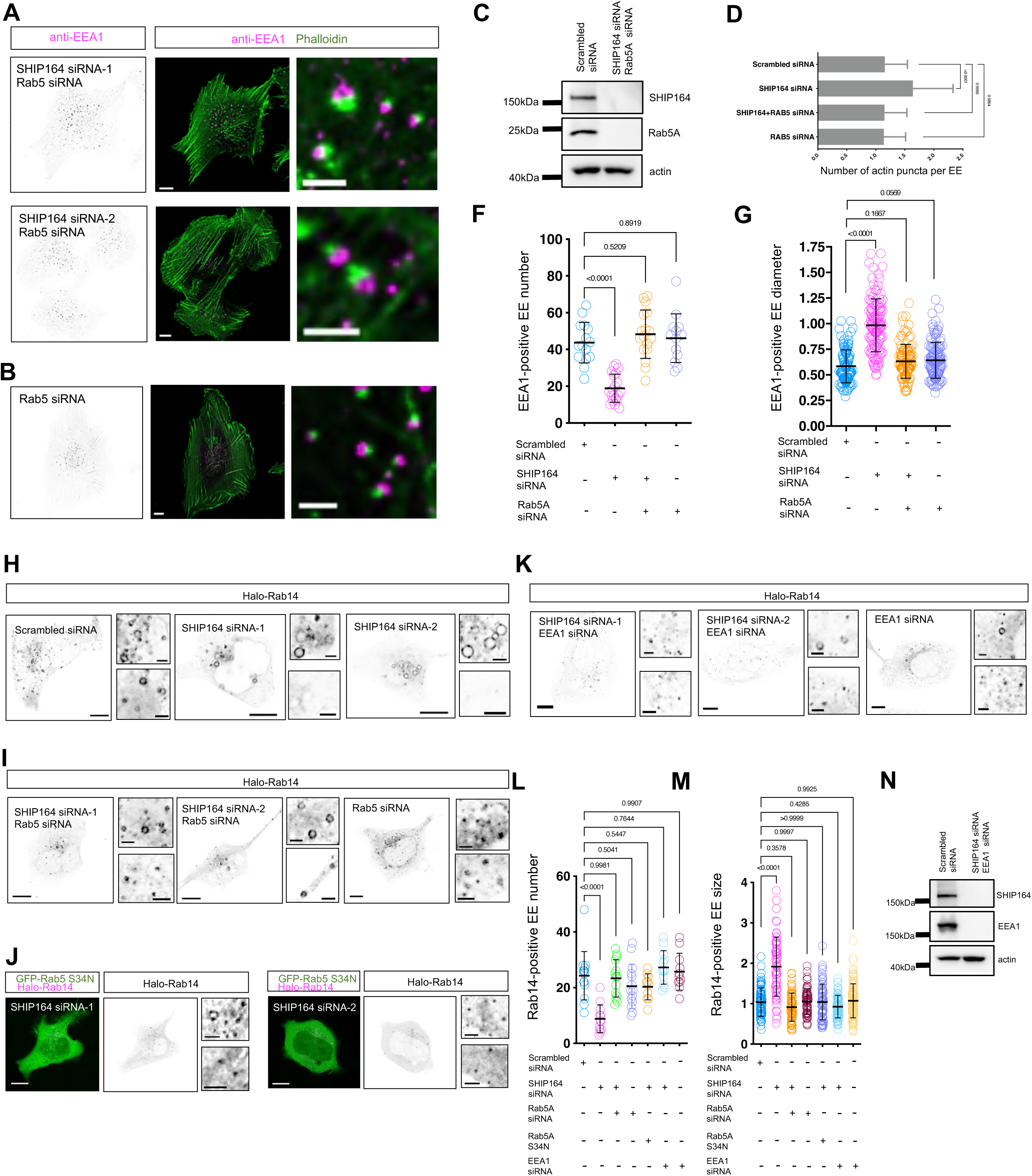
SHIP164 depletion did not substantially affect the number or size of EE. **A, b**. Representative images of fixed RPE1 cells stained with antibodies against endogenous EEA1 (magenta) and phalloidin (green) upon SHIP164 and Rab5 siRNAs or Rab5 siRNA alone with insets. **C.** Immunoblots showing the efficiency of SHIP164 and Rab5 depletion. **D.** The number of actin puncta per EE in scrambled (10 cells) or SHIP164 siRNA (10 cells), SHIP164 and Rab5 siRNA (11 cells), or Rab5 siRNA (10 cells)-treated cells from 3 independent assays. Ordinary one-way ANOVA with Tukey’s multiple comparisons test. Mean ± SD. **F, G**. The number (**F**) or size (**G**) of EEA1-positive EEs per cell in scrambled or SHIP164 siRNA-treated cells. More than 20 cells from 3 independent assays were quantified for each condition. Ordinary one-way ANOVA with Tukey’s multiple comparisons test. Mean± SD. **H, I**. Representative images of live HEK293 cells expressing Halo-Rab14 upon scrambled (left; **H**), SHIP164 (right; **H**), SHIP164 and Rab5 (left; **I**) siRNAs or Rab5 siRNA alone (right; **I**) with two insets. **J.** Representative images of live HEK293 cells expressing dominant-negative Halo-Rab5 S34N and Halo-Rab14 upon SHIP164 depletion with two insets. **K.** Representative images of live HEK293 cells expressing Halo-Rab14 upon SHIP164 and EEA1 (left) or EEA1 siRNA alone with two insets. **L, M**. The number (**L**) or size (**M**) of Rab14-positive EEs per cell in scrambled, SHIP164, SHIP164 and Rab5, Rab5 alone, SHIP164 and EEA1, or EEA1 siRNA alone treated cells. More than 20 cells from 3 independent assays were quantified for each condition. Ordinary one-way ANOVA with Tukey’s multiple comparisons test. Mean ± SD. **N**. Immunoblots showing the efficiency of SHIP164 and EEA1 depletion. Scale bar, 10μm in the whole cell images and 2μm in the insets in (A, B, & H-K).

**Fig. S13.**
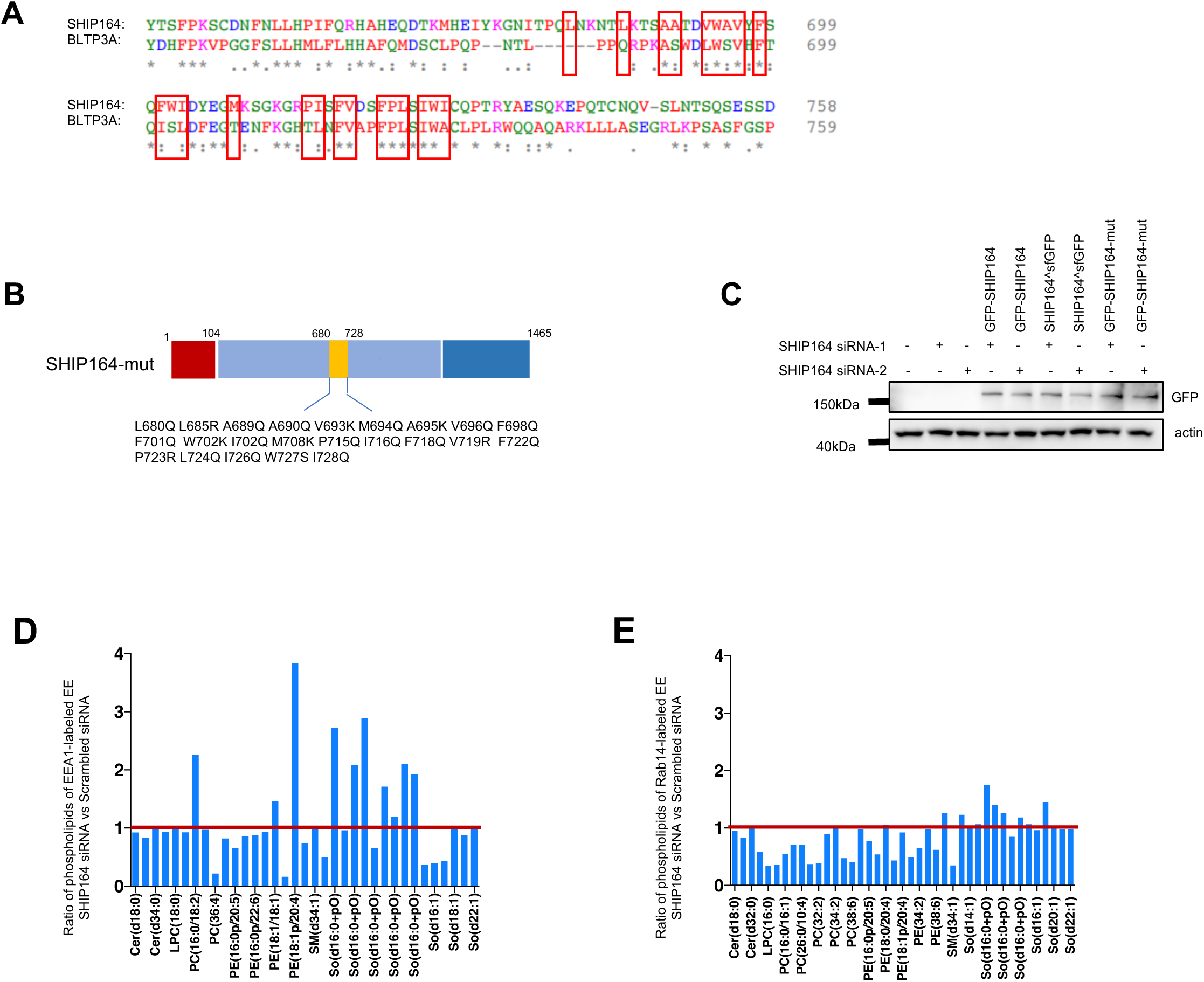
Supplemental results to Figure 4. **A.** Alignment of a region in the middle of lipid transfer domain between SHIP164 and BLTP3A. **B.** Diagram showing the mutated residues in the middle of SHIP164. **C.** Immunoblots showing the level of siRNA-resistant GFP-SHIP164, GFP-SHIP164-mut, or SHIP164^sfGFP in the rescue experiments in Fig.4. **D, E**. Ratio of phospholipid species of GFP-Rab14 (**E**), or GFP-EEA1 (**F**)-labeled EEs in SHIP164-depleted cell normalized to those in scrambled siRNA-treated cells, from three independent assays. Red line denoted ratio=1.

**Fig. S14.**
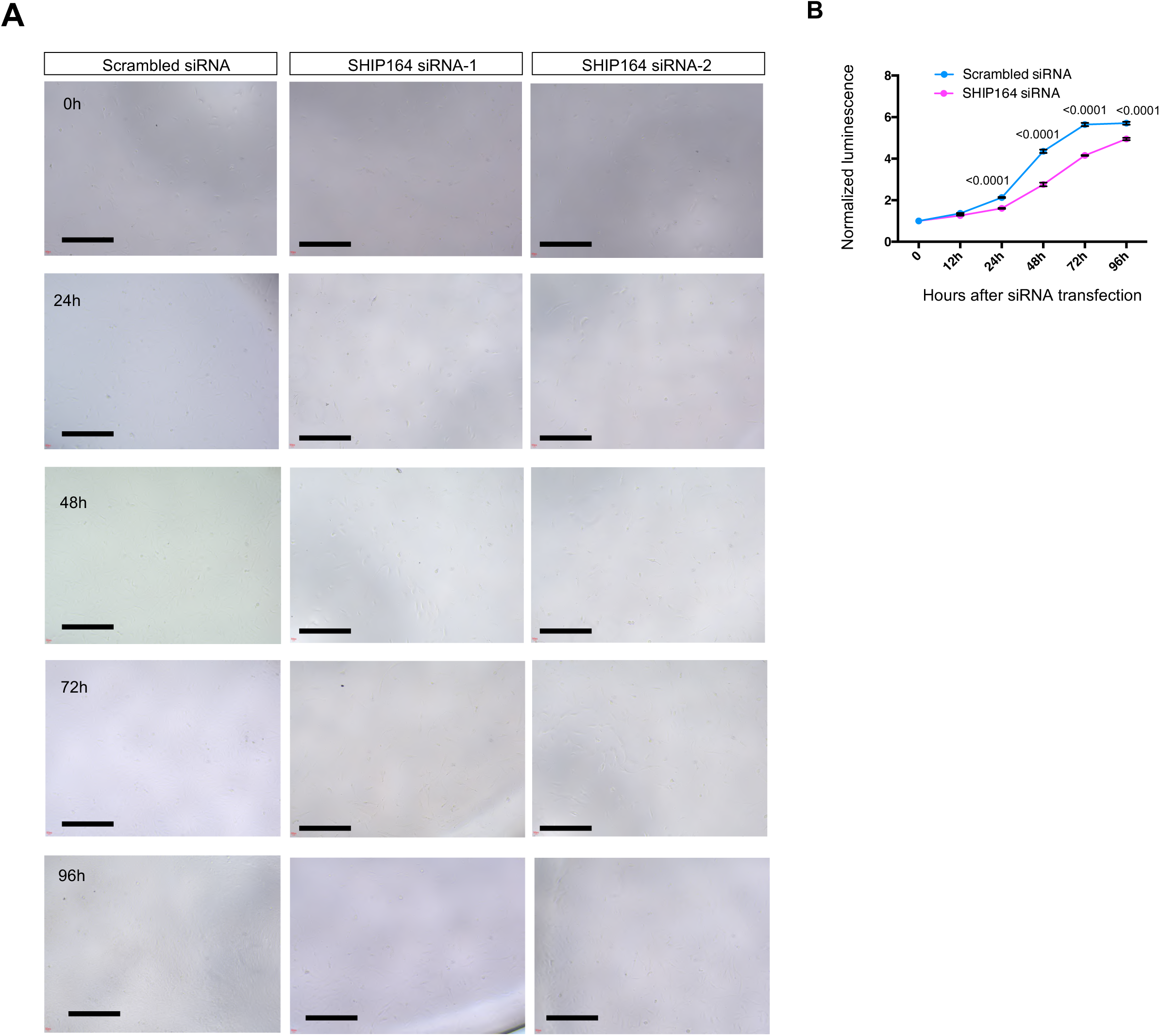
SHIP164 depletion affected cell growth. **A.** Representative bright field images of RPE1 cells at 0h, 12h, 24h, 48h, 72h, or 96h after scrambled or SHIP164 siRNA treatments. **B.** Normalized luminescence of scrambled or SHIP164 siRNA treated cells as in (**A**) from 4 independent experiments. Two-tailed unpaired student t-test. Mean ± SD. Scale bar, 100μm.

**Fig. S15.**
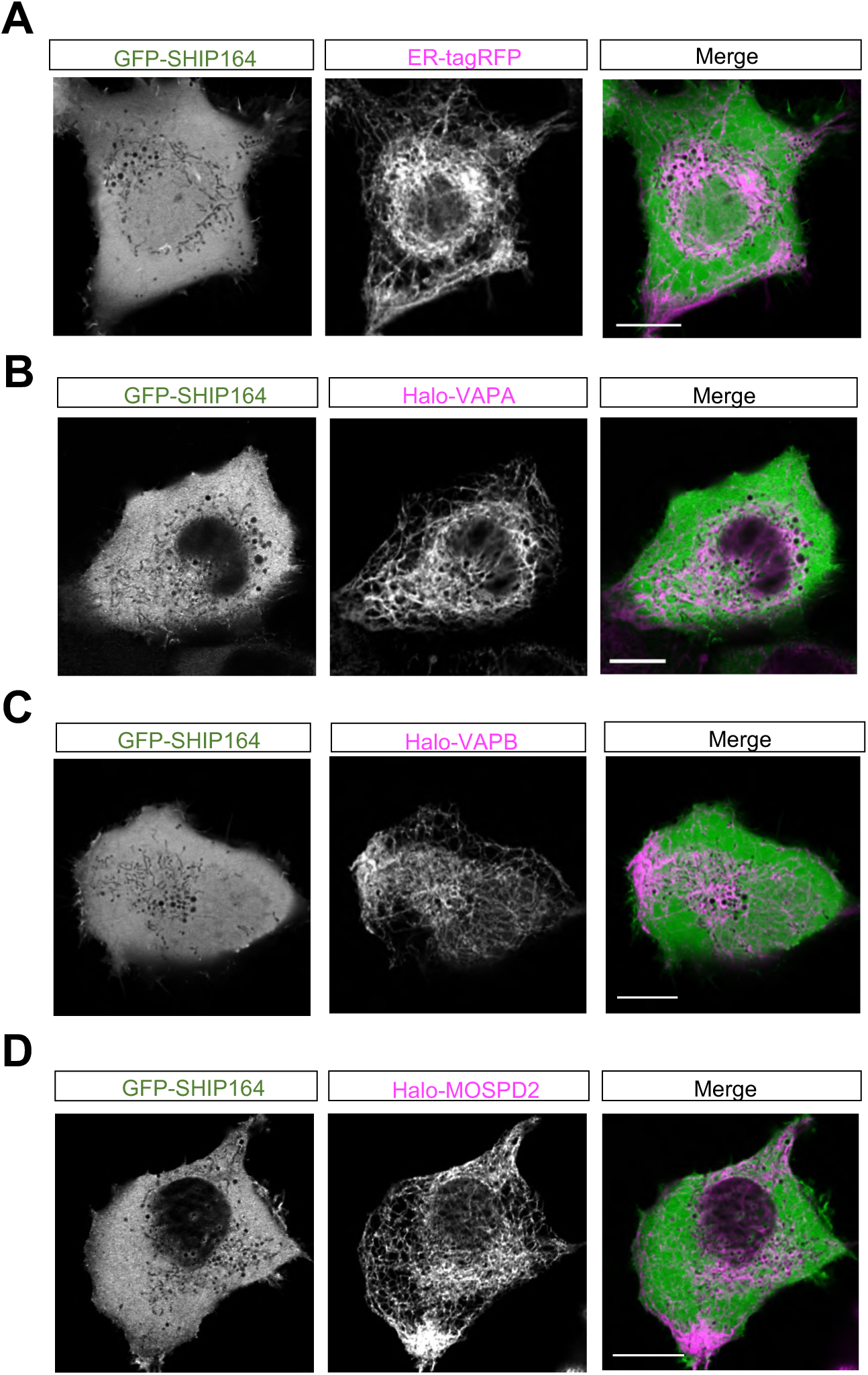
SHIP164 cannot be recruited to the ER by adaptors. **A-D**. Representative images of HEK293 cells expressing either alone ER-tagRFP (magenta; **A**), Halo-VAPA (magenta; **B**), Halo-VAPB (magenta; **C**) or Halo-MOSPD2 (magenta; **D**), along with GFP-SHIP164. Scale bar, 10μm in the whole cell images and 2μm in the insets in (A-D).

**Fig. S16.**
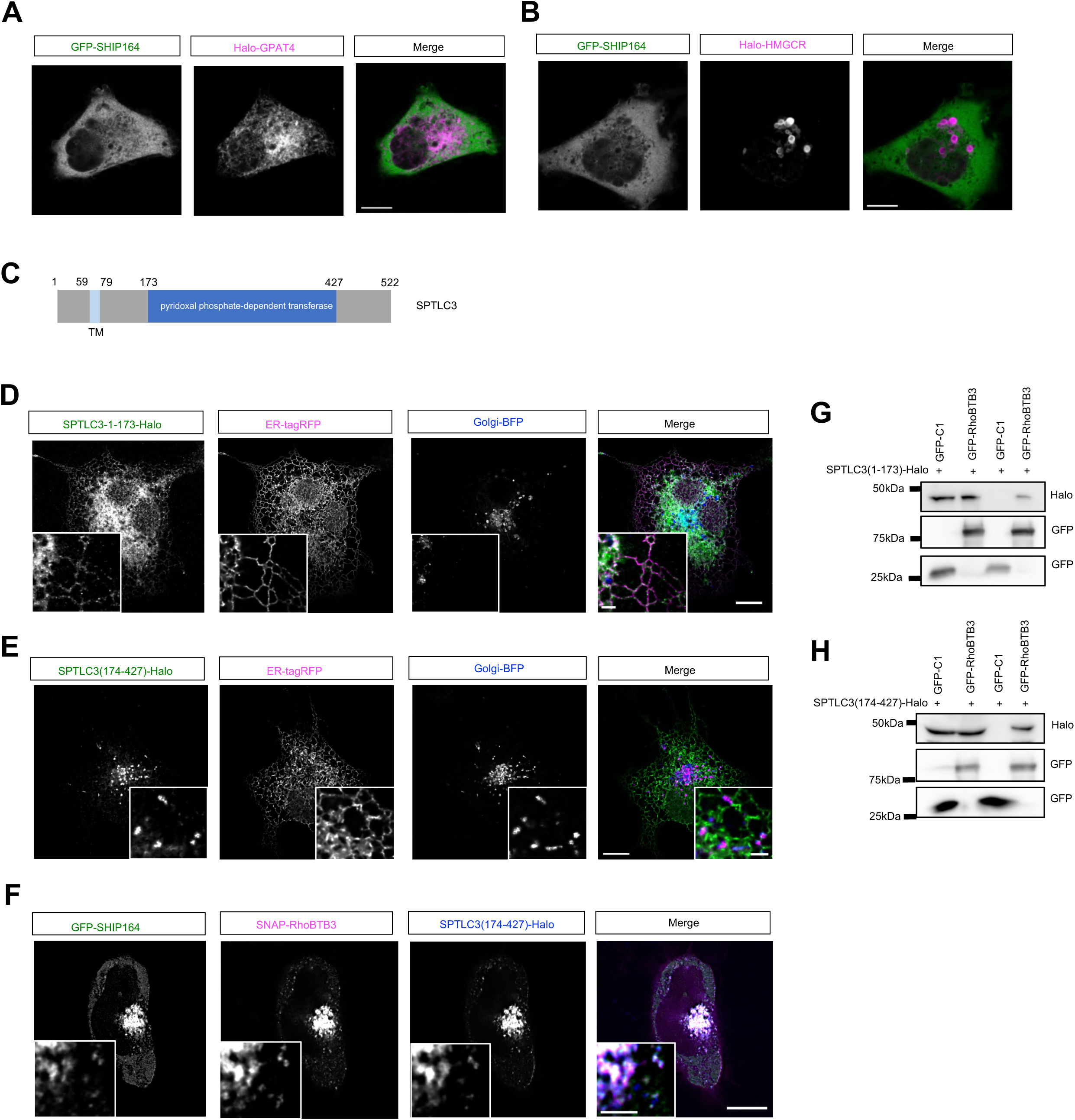
The NT of SPTLC3 targets the ER, while its CT recognizes the Golgi complex. **A, B**. Representative images of HEK293 cells expressing either Halo-GPAT4 (magenta; **A**) or Halo-HMGCR (magenta; **B**) along with GFP-SHIP164 (green). **C.** Domain organization of SPTLC3. **D-E**. Representative images of HEK293 cells expressing either SPTLC3-1-173-Halo (green) or SPTLC3-174-427-Halo (green), along with ER-tagRFP (magenta) and Golgi- BFP with insets. **F**. Representative images of a HEK293 cell expressing GFP-SHIP164 (green), SNAP- RhoBTB3 (magenta), and SPTLC3-174-427-Halo (blue) with insets. **G, H**. GFP-Trap assays demonstrated interactions between GFP-RhoBTB3 and either SPTLC3-1-173-Halo (**E**), or SPTLC3-174-427-Halo (**F**) in HEK293 cells. Scale bar, 10μm in the whole cell images and 2μm in the insets in (A, B, & D-F).

**Fig. S17.**
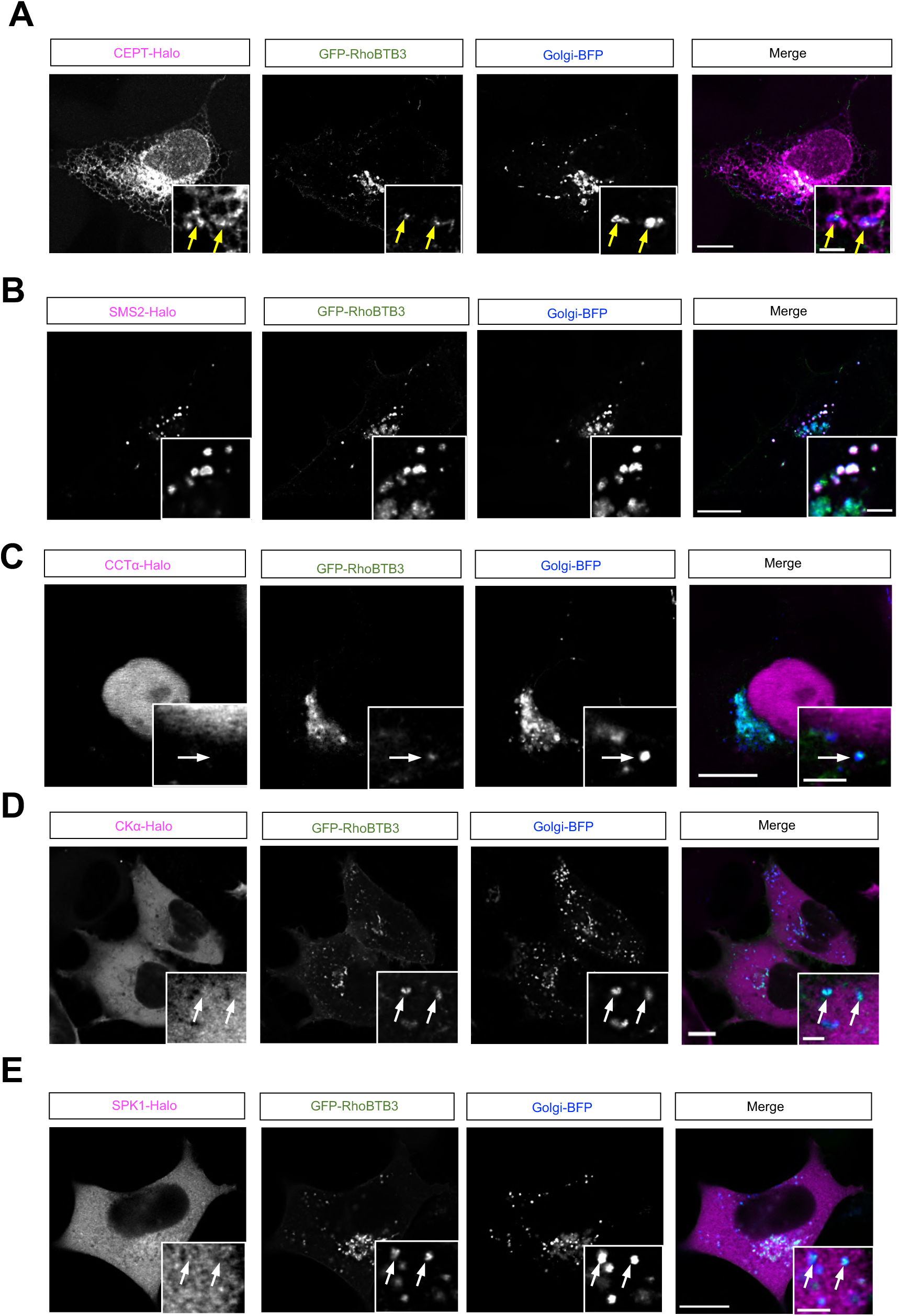
The association between enzymes in phospholipid synthesis pathways and RhoBTB3-positive Golgi. **A-E**. Representative images of HEK293 cells expressing either CEPT-Halo (magenta; **A**), SMS2-Halo (magenta; **B**), CCTα-Halo (magenta; **C**), CKα-Halo (magenta; **D**), SPK1-Halo (magenta; **E**), along with GFP-RhoBTB3 (green) and Golgi-BFP (blue) with insets. Yellow arrows denoted that associations between these enzymes and RhoBTB3, while white arrows indicated RhoBTB3-positive Golgi vesicles not associating with these enzymes. Scale bar, 10μm in the whole cell images and 2μm in the insets in (A-E).

**Fig. S18.**
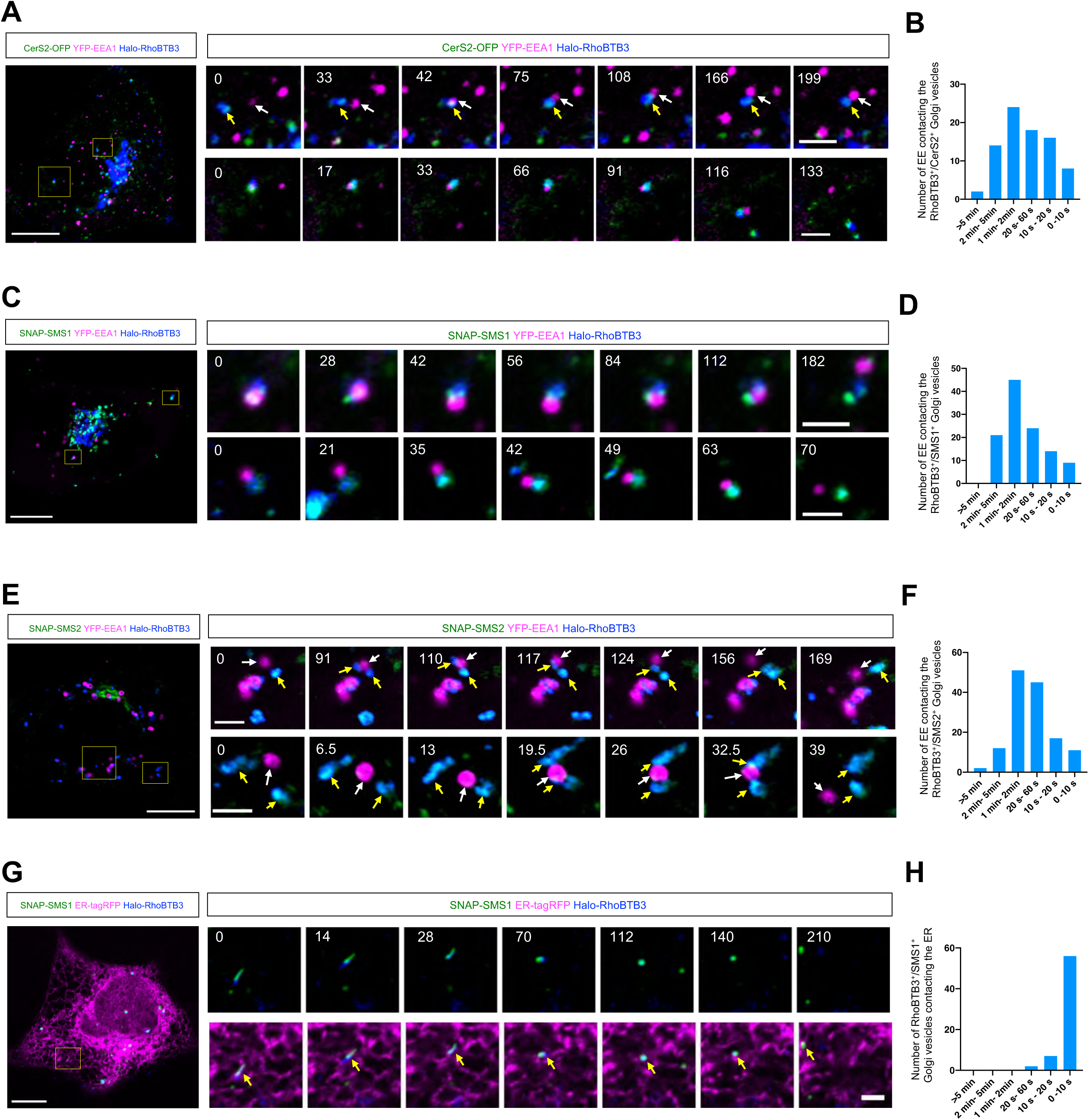
The RhoBTB3 Golgi vesicles frequently contact EEs. **A.** Representative images of a HEK293 cell expressing CerS2-OFP (green), YFP-EEA1 (magenta), and Halo-RhoBTB3 (blue). Left: whole cell images; right: time-lapse images showing dynamic interactions between EEs and RhoBTB3/CerS2-labled Golgi vesicles. **B.** Duration of the contacts between EEA1-labeled EEs and RhoBTB3/CerS2-labled Golgi vesicles from 3 independent assays. **C, E**. Representative images of a HEK293 cell expressing either SNAP-SMS1 (green; **C**), or SNAP-SMS1 (green; **E**), along with YFP-EEA1 (magenta), and Halo-RhoBTB3 (blue). **D, F**. Duration of the contacts between EEs and either RhoBTB3/SMS1 (**D**) or RhoBTB3/SMS2 (**F**) labeled Golgi vesicles from 3 independent assays. **G.** Representative images of a HEK293 cell expressing SNAP-SMS1 (green), ER-tagRFP (magenta), and Halo-RhoBTB3 (blue). **H.** Duration of the contacts between the ER and RhoBTB3/SMS1-labled Golgi vesicles from 3 independent assays. Scale bar, 10μm in the whole cell images and 2μm in the insets in (A, C, E & G).

